# Injectable butyrate-prodrug micelles induce long-acting immune modulation and suppress autoimmune arthritis in mice

**DOI:** 10.1101/2023.08.20.554028

**Authors:** Shijie Cao, Erica Budina, Ruyi Wang, Matthew Sabados, Ani Solanki, Mindy Nguyen, Kevin Hultgren, Arjun Dhar, Jeffrey A. Hubbell

## Abstract

Dysbiosis is linked to autoimmune diseases such as rheumatoid arthritis (RA), where microbial metabolites, such as short chain fatty acids (SCFAs), mediate the so-called gut-joint axis. The therapeutic potential of SCFAs is limited due to the frequent and high oral dosage requirements. RA is characterized by aberrant activation of peripheral T cells and myeloid cells. We aim to deliver butyrate, an SCFA, directly to the lymphatics using a polymeric micelle as a butyrate prodrug, creating a depot for inducing long-lasting immunomodulatory effects. Notably, negatively charged micelles (Neg-ButM) demonstrate superior efficacy in targeting the lymphatics post-subcutaneous administration, and were retained in the draining lymph nodes, spleen, and liver for over a month. In a mouse RA model, we found that Neg-ButM substantially mitigated arthritis symptoms and promoted tolerogenic phenotypes in T cells and myeloid cells, both locally and systemically. These findings suggest potential applications of this approach in treating inflammatory autoimmune diseases.

## Introduction

Rheumatoid arthritis (RA) is an autoimmune disease characterized by chronic inflammation and progressive damage to the synovial joints. It affects approximately 1.3 million adults in the United States, comprising ∼0.5% of the population (*1*). Current treatments for RA often include nonsteroidal anti-inflammatory drugs, disease-modifying anti-rheumatic drugs, and biologics, yet these interventions are often limited by side effects, variable responses, and high cost (*2*). In the quest for more effective RA prevention and treatment, the complex relationship between the gut microbiome and immune responses in the RA disease settings has recently come into focus (*3–6*). Dysbiosis, or the imbalance of commensal bacteria, can affect the onset and progression of RA, with microbial metabolites playing a significant role (*7*, *8*). Among these metabolites, the short-chain fatty acids (SCFAs), including butyrate, are of particular interest due to their potent anti-inflammatory and immunomodulatory effects (*9–11*).

Butyrate, derived from the microbial fermentation of dietary fiber, serves as a key energy source for colonocytes and helps maintain intestinal homeostasis (*9*, *12*). As a histone deacetylase (HDAC) inhibitor, butyrate can modify chromatin structures, regulate gene expression, and facilitate the function of regulatory T cells (Tregs) through upregulation of forkhead box P3 (Foxp3) (*13–15*). Concurrently, it suppresses NFκB activation, inhibits IFNγ production, and amplifies PPARγ expression (*16–19*). Butyrate also exerts anti-inflammatory effects on immune cell compartments through signaling via specific G-protein coupled receptors (GCPRs), such as GPR41, GPR43, and GPR109A (*20–23*). Collectively, due to these properties, butyrate has significant potential as a therapeutic strategy, particularly in the treatment of immunological disorders, including autoimmune diseases. Despite its advantages, the clinical translation of butyrate has been hindered by unpleasant odor and taste, rapid gut metabolism upon oral administration, and difficulties for other routes (e.g. intravenous and rectal administration) intended for chronic management (*24–27*). These challenges underscore the need for innovative delivery methods for butyrate.

In this context, the lymphatic system emerges as a potential target, given its roles in clearing fluid and inflammatory cells from inflamed tissue, and facilitating immune tolerance (*28*, *29*). Recent studies have explored the use of nanocarriers to deliver agents to the lymph nodes (LNs) (*30*, *31*). The lymphatic vessels exhibit wider inter-endothelial junctions than vascular capillaries, allowing larger carriers (10-100 nm) to enter more efficiently from the interstitium (*32*). Building on this idea, here we have engineered butyrate-prodrug micelles to target the LNs and modulate local or systemic immune responses.

Previously, we developed neutral and negatively charged polymeric micelles, termed NtL-ButM and Neg-ButM, respectively (**Fig. 1A, B**) (*11*). Both contain 28 wt% butyrate and have similar sizes of ∼40 nm, making them suitable for LN targeting. We have demonstrated their efficacy through oral administration in treating food allergies and colitis in mice (*11*). In this study, we investigate their potential for RA treatment through subcutaneous (s.c.) administration. Both micelles demonstrated *in vitro* Treg induction and suppression of bone marrow-derived dendritic cells (BMDCs) activated by lipopolysaccharide (LPS). However, *in vivo*, we found that Neg-ButM accumulated more in draining LNs than NtL-ButM, for over a month, primarily being taken up by macrophages and dendritic cells. This leads to significant *in vivo* bioactivity differences when injected s.c. into mice. In antibiotic-treated mice, three weekly doses of Neg-ButM significantly increased the numbers of CD25^+^ Foxp3^+^ CD4^+^ Tregs in draining LNs compared to either PBS or NtL-ButM treatment. Neg-ButM also suppressed the activation of myeloid cells, particularly macrophages, in the draining LNs upon a LPS stimulation. In a mouse model of collagen antibody-induced arthritis (CAIA), administration of only two s.c. injections of Neg-ButM led to a significant reduction in disease progression, associated with both local and systemic increase in Tregs and upregulated immune checkpoints PD-1 and CTLA-4 on both CD4^+^ and CD8^+^ T cells. Neg-ButM also remarkably downregulated co-stimulatory markers, such as CD86, across several myeloid cells in the LNs, and spleen. In addition to its potential application in RA illustrated here, the immune modulation capacity of the Neg-ButM butyrate nanoscale prodrug suggests exploration other inflammatory conditions.

**Figure 1.**
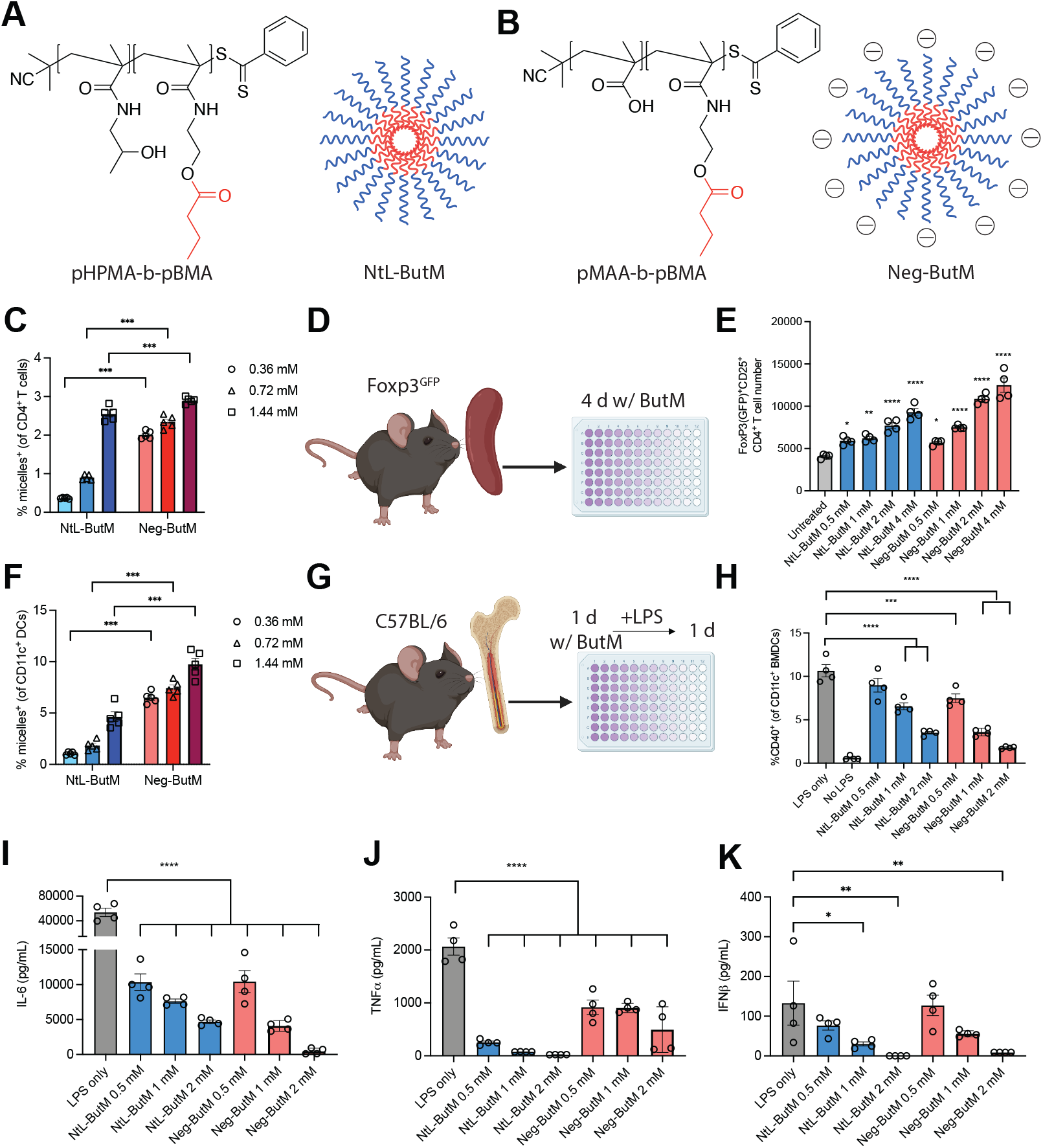
Butyrate micelles induce Tregs and suppress BMDCs in vitro. **A, B**. Chemical structure of pHPMA-b-pBMA (A) or pMAA-b-pBMA (B), which self-assemble into neutral (NtL-ButM) or negatively-charged (Neg-ButM) micelles, respectively. **C.** The proportion of CD4^+^T cells from splenocytes that internalized the fluorescent micelles after 1-hr incubation with the micelles, at micelle concentrations corresponding to butyrate concentrations of 0.36 mM, 0.72 mM, and 1.44 mM. **D-E.** In vitro Treg generation by butyrate micelles. Number of CD25^+^Foxp3^+^CD4^+^ Tregs cells from splenocytes isolated from Foxp3GFP reporter mice and treated with NtL-ButM or Neg-ButM for 4 days. **F.** The proportion of CD11c^+^ DCs from splenocytes that internalized the fluorescent micelles after a 1-hr incubation with the micelles. **G-K**, The proportion of BMDCs expressing CD40 (H), and the level of IL-6 (I), TNFα (J), and IFNβ (K) in the supernatant after incubation with butyrate micelles for 24 hr, followed by the addition of LPS for an additional 18 hr. Data are presented as mean ± SEM. Statistical analyses in E, H, I-K, were performed using a one-way ANOVA with Dunnett’s test. *n* = 4, and each experiment was repeated at least twice. Statistical analyses in C, F were performed using a two-way ANOVA with Bonferroni’s test. *p<0.05, **p<0.01, ***p<0.001, ****p<0.0001. Fig. 1D, G were created with BioRender.com.

## Results

### Butyrate micelles induce Tregs and suppress dendritic cell activity *in vitro*

Gut microbiome-derived butyrate is known to induce Treg differentiation from naïve CD4^+^ T cells in the periphery (*17–19*). Butyrate can achieve this through HDAC inhibition and by influencing the behavior of dendritic cells (DCs) (*20*, *21*). DCs can modulate their cell-surface checkpoint molecules and secrete tolerance-inducing cytokines like cytokines TGFβ that promote Treg generation (*33*). In addition, butyrate’s suppression of inflammatory cytokines such as IL-12, IL-6, and tumor necrosis factor-α (TNFα) may further facilitate Treg generation (*34*, *35*).

We hypothesized that our butyrate micelles, by providing a direct interaction between butyrate and key immune cells such as T cells and DCs, would promote Treg generation. We started by conducting an *in vitro* cell uptake study. After incubating isolated splenocytes from C57BL/6 mice with fluorescently labeled butyrate micelles and analyzing them using flow cytometry, we found that the Neg-ButM micelles were taken up more readily by CD4^+^ T cells than the NtL-ButM micelles at equal concentrations (**Fig. 1C**). This uptake increased proportionally with the concentration of micelles.

To evaluate the regulatory efficacy of the butyrate micelles *in vitro*, we used Foxp3^GFP^ reporter mice, which have green fluorescent protein (GFP) attached to the protein Foxp3 (**Fig. 1D**). Splenocytes from these mice were cultured for 4 d with different concentrations of either Neg-ButM or NtL-ButM, with an untreated control as the reference (**Fig. 1E**). Both micelles induced Treg generation over the four-day period, but Neg-ButM were more potent at the same loaded butyrate concentration, increasing Treg numbers up to 3-fold.

In the *in vitro* cell uptake study, we also assessed the micelle uptake by DCs. Similar to CD4^+^ T cells, Neg-ButM micelles were taken up more significantly by DCs compared to NtL-ButM at the same concentration (**Fig. 1F**). Furthermore, up to 10% of the total DC population exhibited potentially active endocytosis, compared to the nearly 3% of CD4+ T cells that may be physically associated with the micelles.

We then analyzed the potential effect of ButM on CD40 expression on DCs. High levels of CD40 are associated with increased activation of DCs, driving inflammatory responses. On the contrary, lower levels of CD40 can lead to Treg generation and increased tolerance (*36*). We isolated BMDCs from C57BL/6 mice and incubated them with different concentrations of micelles. Following this, lipopolysaccharide (LPS) was added to stimulate the cells, and flow cytometry was used to determine CD40 expression levels (**Fig. 1G**). Both types of micelles were found to suppress CD40 expression, up to a 6-fold decrease from Neg-ButM, compared to cells treated only with LPS (**Fig. 1H**), and the effects were dose-dependent. This aligns well with the Treg induction study and indicates that a portion of the Treg induction by micelles could be attributed to the suppression of DC activation signals.

Cytokine release is a critical function of DCs that influences T cell fate. Overproduction of inflammatory cytokines, particularly by activated DCs, can hinder Treg induction and potentially contribute to autoimmune diseases, as well as promote inflammation (*37*). To determine the impact of butyrate micelles on cytokine production, we measured cytokine levels in the supernatant after incubating BMDCs with butyrate micelles and stimulating them with LPS. Both Neg-ButM and NtL-ButM suppressed the production of the inflammatory cytokines IL-6, TNFα, and IFNβ in a dose-dependent manner (**Fig. 1I-K**). Interestingly, while both micelles performed similarly in terms of overall suppression, NtL-ButM demonstrated superior efficacy in suppressing TNFα. Importantly, this suppressive effect was attributed to the bioactivity of the micelles, not cytotoxicity, as the majority of the cells remained viable after treatment (**Fig. S4B**).

### Lymph node targeting and retention is enhanced by negatively charged butyrate micelles

While both butyrate micelle formulations, Neg-ButM and NtL-ButM, displayed comparable bioactivities *in vitro* regarding Treg induction and DC inhibition, their biodistribution *in vivo*, which could impact their functionality, may differ. Nanoparticles are known to preferably traffic to lymph nodes (LNs) upon injection into the interstitium, utilizing the lymphatic vessels’ wider inter-endothelial junctions compared to vascular capillaries (*32*). Moreover, neutral or positively charged vehicles tend to be trapped in the negatively-charged extracellular matrix of the interstitium (*38*). Prior studies have highlighted that negatively charged nanoparticles might demonstrate superior LN targeting after s.c. administration (*31*, *39*). Therefore, in this study, we sought to compare the biodistribution of the two butyrate micelle formulations and evaluate their lymphatic targeting abilities.

To monitor the *in vivo* biodistribution of the micelles, we s.c. administered fluorescently labeled NtL-ButM or Neg-ButM into the mice’s abdomens (**Fig. 2A**). At specified intervals after injection, we collected major organs and examined the fluorescence signals using an In Vivo Imaging System (IVIS). A striking observation was that the Neg-ButM not only accumulated in the draining inguinal LNs, but were also retained there for an extended period or more than 35 days. This level of accumulation and retention was not witnessed for NtL-ButM (**Fig. 2B**). Further examination revealed that a single injection of Neg-ButM led to significantly higher signal levels in the draining LNs across all the sampled time points. Additionally, Neg-ButM not only localized to the draining LNs but also reached non-draining LNs, including the contralateral inguinal LNs and gut-draining LNs, albeit with less fluorescence signal. Interestingly, this s.c. administration also led to systemic biodistribution. While we did not detect any signal in the blood, possibly due to the limit of detection from diluted fluorescence signals, we identified considerable signal amounts in the spleen and kidney, with the liver showing the highest levels. The signals peaked on the 7th day but remained robust even on the 35th day (**Fig. 2B**).

**Figure 2.**
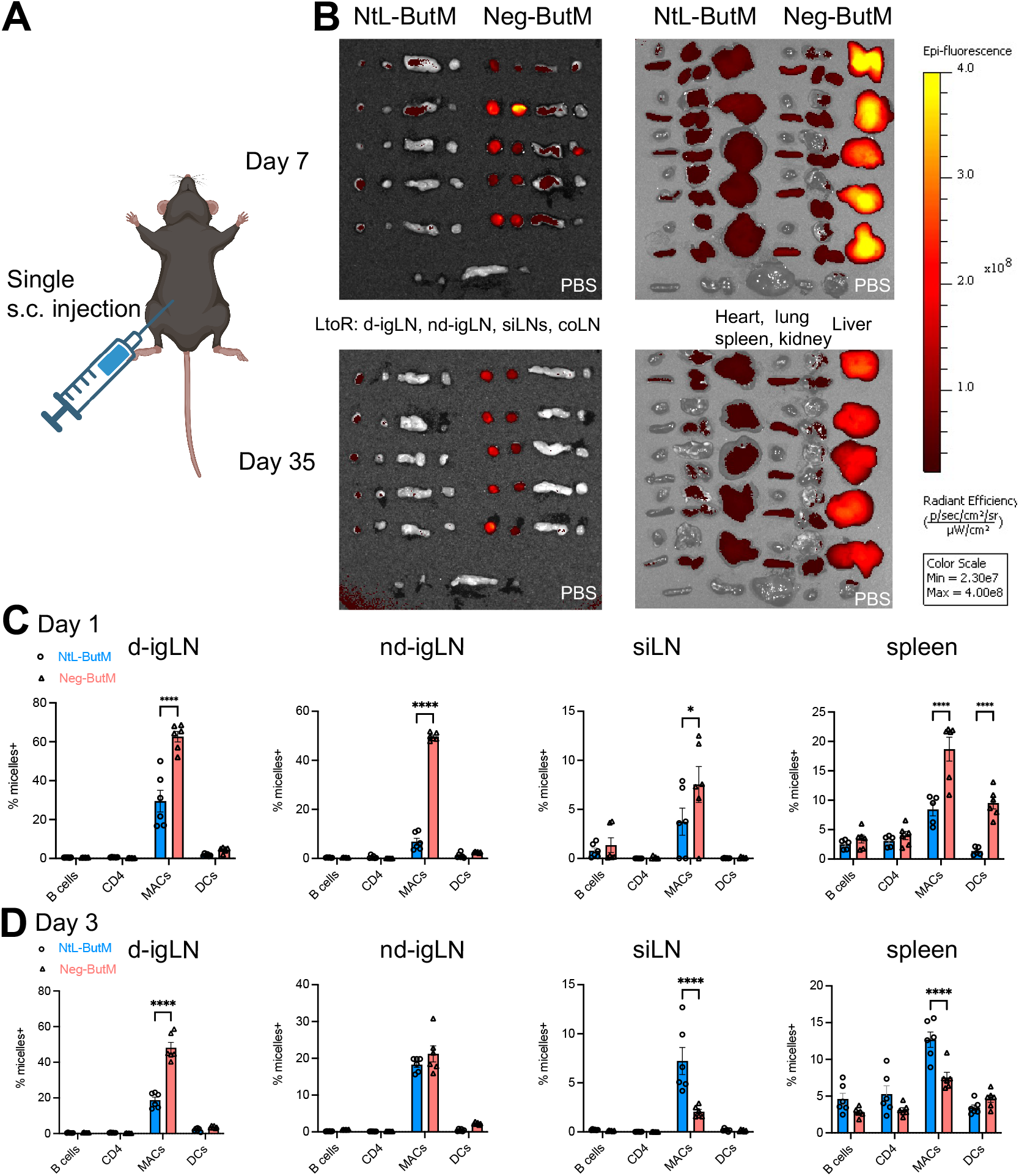
In vivo biodistribution of butyrate micelles. **A, B.** IVIS images of lymph nodes (LNs) and major organs harvested at different time points after mice were subcutaneously injected in the right lower abdomen with fluorescently-labeled NtL-ButM or Neg-ButM. Within a set of LNs from one mouse, left to right: draining and non-draining inguinal LNs (d-igLN, nd-igLN), small intestine LNs (siLNs), and colon LNs (coLN). *n* = 5. **C, D,** The percentage of fluorescently-labeled ButM in different subsets of cells in the d-igLN, nd-igLNs, siLN, or spleen at 1 d (C) or 3 d (D) after mice were subcutaneously injected with micelles. *n* = 6. MAC, macrophages. DC, dendritic cells. Data are presented as mean ± SEM. Statistical analyses were performed using a two-way ANOVA with Bonferroni’s test. *p<0.05, **p<0.01, ***p<0.001, ****p<0.0001.

To further delineate the biodistribution at a cellular level, we digested LNs and spleen from mice on d 1 and d 3 post-injection, obtaining single cell suspensions. Using flow cytometry, we analyzed fluorescently labeled micelle-positive cells across different cell populations. Both micelle forms were predominantly taken up by macrophages in the LNs, with Neg-ButM being assimilated by a larger number of cells compared to NtL-ButM on d 1 (**Fig. 2C, D**). Notably, in the spleen, in addition to macrophages, dendritic cells also took up the micelles, particularly Neg-ButM, which accounted for up to 10% of the cell population. Comparing the two time points, we observed a decrease in cellular uptake from d 1 to d 3 for Neg-ButM.

### Butyrate micelles induce Tregs and suppress macrophages activity *in vivo*

Microbiome-derived SCFAs, especially butyrate, are recognized for fostering the extrathymic differentiation of Tregs (*17*, *18*). As a potential therapeutic candidate, butyrate’s capability to induce gut-localized Tregs has received considerable attention(*17–19*). Peripherally-derived Tregs are induced most efficiently in the LNs, yet current therapies do not efficiently target the LN. We have successfully shown that our butyrate micelles, Neg-ButM and NtL-ButM, are able to induce regulatory T cells *in vitro*. However, *in vivo* biodistribution studies illustrated that Neg-ButM surpasses NtL-ButM in LN targeting after s.c. administration. Consequently, we sought to explore the potential of Neg-ButM in inducing Tregs in draining LNs through s.c. administration.

We utilized the C57BL/6 Foxp3^GFP+^ mice to assess the potential of *in vivo* Treg induction. To delineate the effect of administered butyrate, mice were given antibiotic water throughout the study, thereby eliminating the influence of endogenous butyrate from the gut microbiome. We s.c. administered PBS, NtL-ButM, or Neg-ButM to the mice once a week for three weeks. One week after the final dose, we euthanized the mice and isolated cells from the draining LNs to analyze the Treg population via flow cytometry (**Fig. 3A**). We showed that Neg-ButM administration significantly increased the Foxp3^+^CD25^+^ regulatory T cells by 1.8-fold in the draining LNs, whereas neither NaBut nor NtL-ButM led to Treg increases compared to PBS-treated mice. Thus, despite both butyrate formulations’ ability to induce Tregs *in vitro*, effective *in vivo* induction necessitates LN delivery and prolonged presence, provided by Neg-ButM.

**Figure 3.**
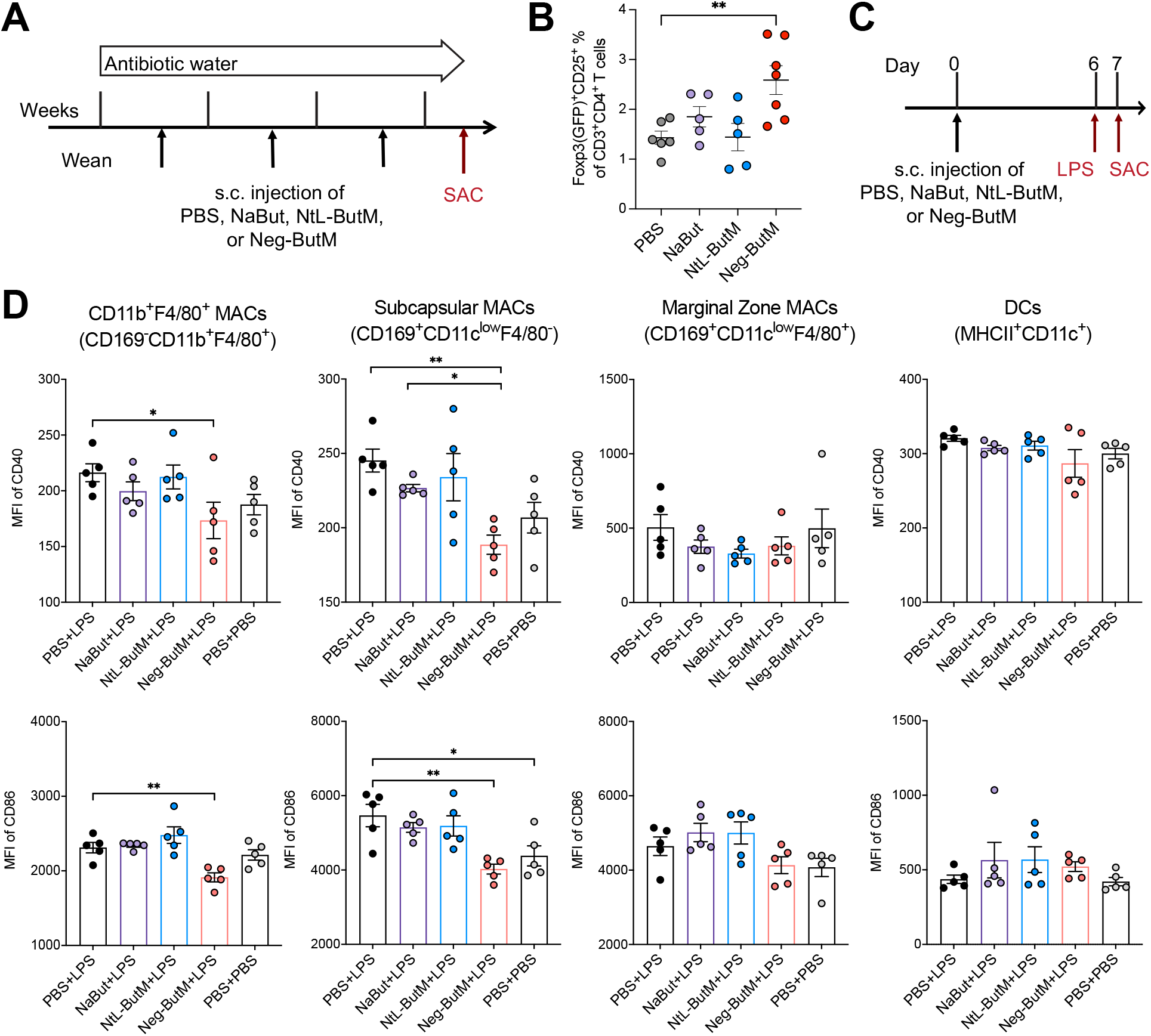
Negatively charged butyrate micelles induce Tregs and suppress LPS-induced activation *in vivo*. **A.** Foxp3^GFP^reporter mice were subcutaneously (s.c) injected with butyrate formulations (equivalent to 1.5 mg of butyrate per mouse) once a week for 3 wk. Mice were provided with antibiotic in the drinking water to eliminate endogenous butyrate. **B.** The percentage of Foxp3^+^CD25^+^ Tregs of CD3^+^CD4^+^ T cells in the draining inguinal LNs. *n* = 5-6. Data were pooled from two experiments. **C.** C57BL/6 mice were s.c. injected with butyrate formulations on day 0, followed by s.c. stimulation with LPS on day 6, and were sacrificed on day 7. **D.** The mean fluorescence intensity (MFI) of CD40 or CD86 expression on CD11b^+^F4/80^+^ macrophages, subcapsular macrophages (CD169^+^ CD11c^low^F4/80^-^), marginal zone macrophages (CD169^+^ CD11c^low^F4/80^+^), and CD11b^+^ dendritic cells in the draining LNs. *n* = 5. Data are presented as mean ± SEM. Statistical analyses were performed using a one-way ANOVA with Dunnett’s test. *p<0.05, **p<0.01, ***p<0.001, ****p<0.0001.

We further evaluated the ability of LN-targeted Neg-ButM to suppress the activation of myeloid cells. Using a mouse model of subcutaneous LPS stimulation, we assessed activation markers, including CD40 and CD86, on the major myeloid cells in the draining LNs. Mice were s.c. injected with either PBS, NaBut, NtL-ButM, or Neg-ButM at the abdominal site. On day 6, mice were challenged with LPS at the same injection site and euthanized the next day for cellular analysis.

The Neg-ButM treatment significantly inhibited CD40 and CD86 expression on CD169^-^ CD11b^+^F4/80^+^ macrophages and CD169^+^ CD11c^low^F4/80^-^ subcapsular macrophage in the draining LNs post-LPS stimulation (**Fig. 3D**). The macrophages were the primary uptakers of Neg-ButM, as determined by the previous cellular biodistribution study. Interestingly, no significant effect was observed on DCs, likely attributable to their limited interactions with the micelles, a phenomenon noted in our earlier cellular biodistribution investigations (**Fig. 2C, D**). In contrast to the marked impact of Neg-ButM, neither NtL-ButM nor NaBut demonstrated significant suppression on any of the myeloid cell populations. This finding underscores the crucial need to target prolonged butyrate delivery to the LNs for the optimal manifestation of its bioactivity.

### S.c. injected Neg-ButM mitigates collagen antibody-induced arthritis (CAIA) in mice

Emerging research suggests that the gut microbiota composition significantly influences the onset and progression of RA (*5*, *7*, *40*). Earlier studies indicated the therapeutic potential of butyrate in alleviating RA symptoms by interacting with key immune cells such as osteoclasts and T cells (*41*). Given the enhanced accumulation of Neg-ButM in LNs and its specific bioactivity in inducing Tregs and suppressing myeloid cells upon s.c. administration, we aimed to evaluate its efficacy in the collagen antibody-induced arthritis (CAIA) mouse model of RA. This model imitates the disease through passive immunization using an anti-collagen antibody cocktail, followed by LPS administration, thereby instigating a sequence of innate immune cell infiltration into the joints, leading to inflammation and swelling within a week post-immunization (*42*).

Our experimental design involved two different injection sites for s.c. administration. Initially, we sought to directly target the site of inflammation in the induced arthritis, opting for injections in the four hocks of the mice. We hypothesized that this approach would influence the lymphatic cells in the hock-draining LNs, inhibiting inflammation in the paw joints directly. Mice were pre-treated with either PBS or Neg-ButM at low (2.5 mg/mouse) or high (10 mg/mouse) doses twice, one week apart, before the induction of arthritis (**Fig. S6A**). Notably, Neg-ButM treatment significantly reduced the development of arthritis in treated mice compared to those treated with PBS (**Fig. S6B, C**). Upon homogenizing the harvested paws, we detected lower levels of the inflammatory cytokine IL-6 in mice treated with Neg-ButM, compared to the PBS-treated group (**Fig. S6D**). When analyzing the disease progression clinical score and cytokines in the paws, we observed no significant differences between the two Neg-ButM dosages.

We further characterized the immune cells in the hock-draining LNs and spleen. In both tissues, Neg-ButM treatment resulted in increased Tregs and enhanced PD-1 expression on both CD4^+^ and CD8^+^ T cells (**Fig. S6E-H**). While a dose-dependent effect from Neg-ButM was not evident in the hock-draining LNs, the high dose of Neg-ButM led to a substantial increase in the frequency of Tregs and PD-1^+^ cells among CD4^+^ and CD8^+^ T cells in the spleen. As to myeloid cells in the draining LNs, we also observed a dose-dependent effect of Neg-ButM in suppressing CD86, CD40, and MHCII expression on macrophages in the draining LNs (**Fig. S7A**). Intriguingly, Neg-ButM treatment increased the frequency of CD11b^+^Ly6C^+^Ly6G^-^ cells, known as myeloid-derived suppressor cells (MDSCs) (*43*, *44*), in a dose-dependent manner and suppressed MHCII expression on these cells (**Fig. S7B**). In DCs (CD11c^+^ and CD11b^+^CD11c^+^ cells), Neg-ButM decreased CD40 and CD86 expression, indicating a broad suppression effect of butyrate micelles on these myeloid cells (**Fig. S7C**).

Because of these dose-dependent effects observed in the cellular readout, we elected to continue further investigation with the high dose regimen. We returned to the abdominal site of s.c. administration, adhering to precedent studies’ methods and acknowledging the potential for better clinical relevance. We also conducted further measurements to assess the disease progression and the impact of Neg-ButM s.c. treatment (**Fig. 4A**).

**Figure 4.**
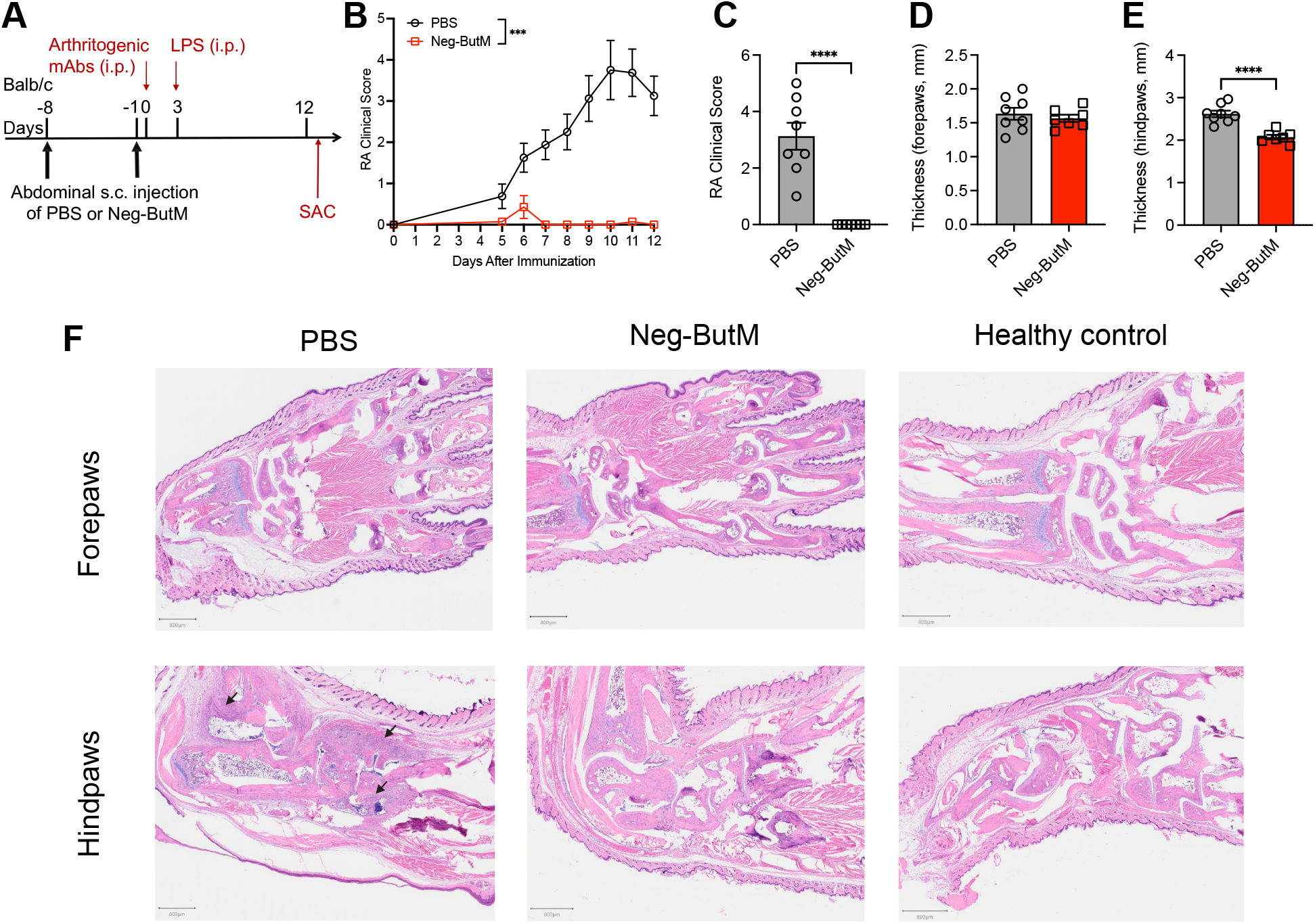
Negatively charged butyrate micelles suppress arthritis progression. **A.** Experimental scheme of the collagen antibody-induced arthritis (CAIA) model. Mice were subcutaneously injected with PBS (*n* = 8) or Neg-ButM (10 mg) (*n* =7) twice on day -8 and day -1. CAIA was induced by passive immunization with an anti-collagen antibody cocktail on day 0, followed by intraperitoneal injection of lipopolysaccharide (LPS). **B.** Arthritis scores were measured daily after the immunization. The areas under the curve of clinical scores were compared. **C.** Arthritis scores of PBS or Neg-ButM treated mice on day 12. **D, E.** The thickness of forepaws (D) or hindpaws (E) measured on day 12 from mice treated with PBS or Neg-ButM. **F.** Representative images of mouse joints from paws stained with hematoxylin and eosin on day 12 in each treatment group. The representative mouse in the PBS group, as shown in the image, had a total clinical score of 4. All mice in the Neg-ButM treated group and the healthy control had a clinical score of 0. Bar = 800 μm. Arrows indicate immune cell infiltration. Data are presented as mean ± SEM. Statistical analyses were performed using Student’s t test. ***p<0.001, ****p<0.0001.

In this experiment, we observed that two abdominal s.c. injections of Neg-ButM effectively suppressed disease progression post-CAIA induction. All mice treated with Neg-ButM showed an arthritis clinical score of zero, contrasting with PBS-treated mice which all exhibited swelling and inflammation in at least one paw (**Fig. 4B-E**). We noted that the most severe inflammation occurred in the hindpaws, not the forepaws, hence significant differences in paw thickness were only apparent in the hindpaws. Histological images revealed immune cell infiltration in the hindpaws of PBS-treated mice, a phenomenon absent in Neg-ButM treated mice (**Fig. 4F, Fig. S10**). Consequently, our findings suggest that abdominal s.c. administration of Neg-ButM yields comparable suppression of arthritis progression in the CAIA model as direct s.c. injection to the hocks.

### S.c. injected Neg-ButM in the abdomen modulates T cells and myeloid cells in lymphoid organs

The successful suppression of disease progression in CAIA mice following two abdominal s.c. injections of Neg-ButM compelled us to delve deeper into understanding how this treatment affects the immune cell characterization in various LNs and spleen. After sacrificing the mice, we examined the inguinal LNs (which are the injection-site draining LNs), the hock-draining LNs (including pooled popliteal, axillary, and brachial LNs) that are disease-relevant LNs, and the spleen, which serves as a circulatory hub for immune cells and is indicative of systemic immune modulation.

We observed that Neg-ButM treatment substantially increased the number of regulatory CD4^+^ T cells in both the inguinal and hock-draining LNs, as well as in the spleen. This was demonstrated by the elevated percentage of Foxp3^+^CD25^+^ cells within the total CD4^+^ T cell population (**Fig. 5A-C**). These findings align with our prior *in vivo* Treg induction study (**Fig. 3B**) and observations in CAIA mice treated with Neg-ButM in the hocks (**Fig. S6E, H**). Simultaneously, we observed a significant increase in the expression of PD-1 and CTLA-4 on both CD4^+^ and CD8^+^ T cells across all LNs and the spleen. This increase was particularly noticeable in the spleen, where we observed a nearly 1.6-fold increase in the PD-1^+^ and CTLA-4^+^ CD4^+^ T cells, and a close to 4-fold increase of PD-1^+^ among CD8^+^ T cells (**Fig. 5C**). These inhibitory checkpoint cellular markers contribute to the suppression of T-cell activation, thus reinforcing immune tolerance and control.

**Figure 5.**
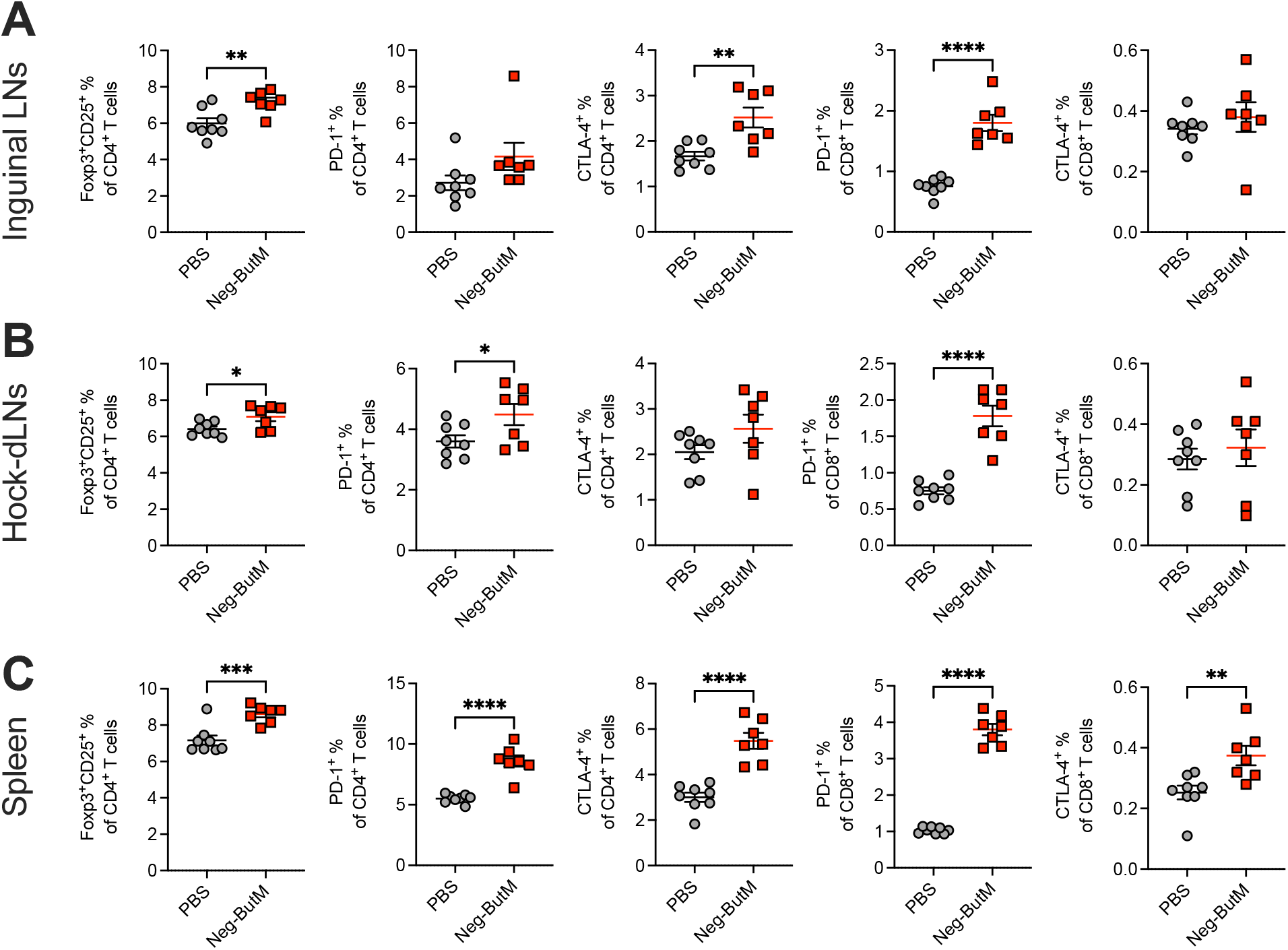
Negatively charged butyrate micelles modulate T cell responses in arthritis. The percentage of Foxp3^+^CD25^+^ regulatory T cells, PD-1^+^, CTLA-4^+^ cells of CD3^+^CD4^+^ T cells, or PD-1^+^, CTLA-4^+^ cells of CD3^+^CD8^+^ T cells in the inguinal lymph nodes (LNs) (**A**), hock-draining LNs (**B**), or spleen (**C**), from mice treated with PBS or Neg-ButM in Fig. 4. Data are presented as mean ± SEM. Statistical analyses were performed using Student’s t test. ***p<0.001, ****p<0.0001.

Myeloid cells such as DCs and macrophages are essential to antigen presentation and T cell activation. Specifically, the interaction between CD86 on myeloid cells and CD28 on T cells triggers their activation and proliferation (*45*, *46*). In contrast, CTLA-4, which is predominantly found on T cells and notably on Tregs, competes with CD28 for CD86 binding, thereby transmitting an inhibitory signal to T cells(*47*). Given our observation of a significant increase of CTLA-4^+^ T cells following Neg-ButM treatment in the CAIA model, we further examined the influence of Neg-ButM on myeloid cells by analyzing CD86 expression across different myeloid cell types in the inguinal LNs, hock-draining LNs, and spleen. Across all the tissues, we noticed a significant downregulation of CD86 on the macrophages, including CD11b^+^CD11c^-^ cells and CD11b^+^F4/80^+^ cells (**Fig. 6A-C**). Similarly, we observed a decrease of CD86^+^ on CD11b^+^Ly6C^+^Ly6G^-^ MDSCs in hock-draining LNs and the spleen (**Fig. 6B-C**), consistent with our findings from direct hock s.c. injection of Neg-ButM (**Fig. S7B**). Moreover, we noted a significant reduction of CD86^+^ on DCs, including both subsets of CD11c^+^ and CD11c^+^CD11b^+^ DCs, across most tissues, except for the CD11c^+^ cells located in the inguinal LNs. Notably, the greatest differences were observed in the spleen, where we documented an approximate 2-fold decrease of CD86 on DCs.

**Figure 6.**
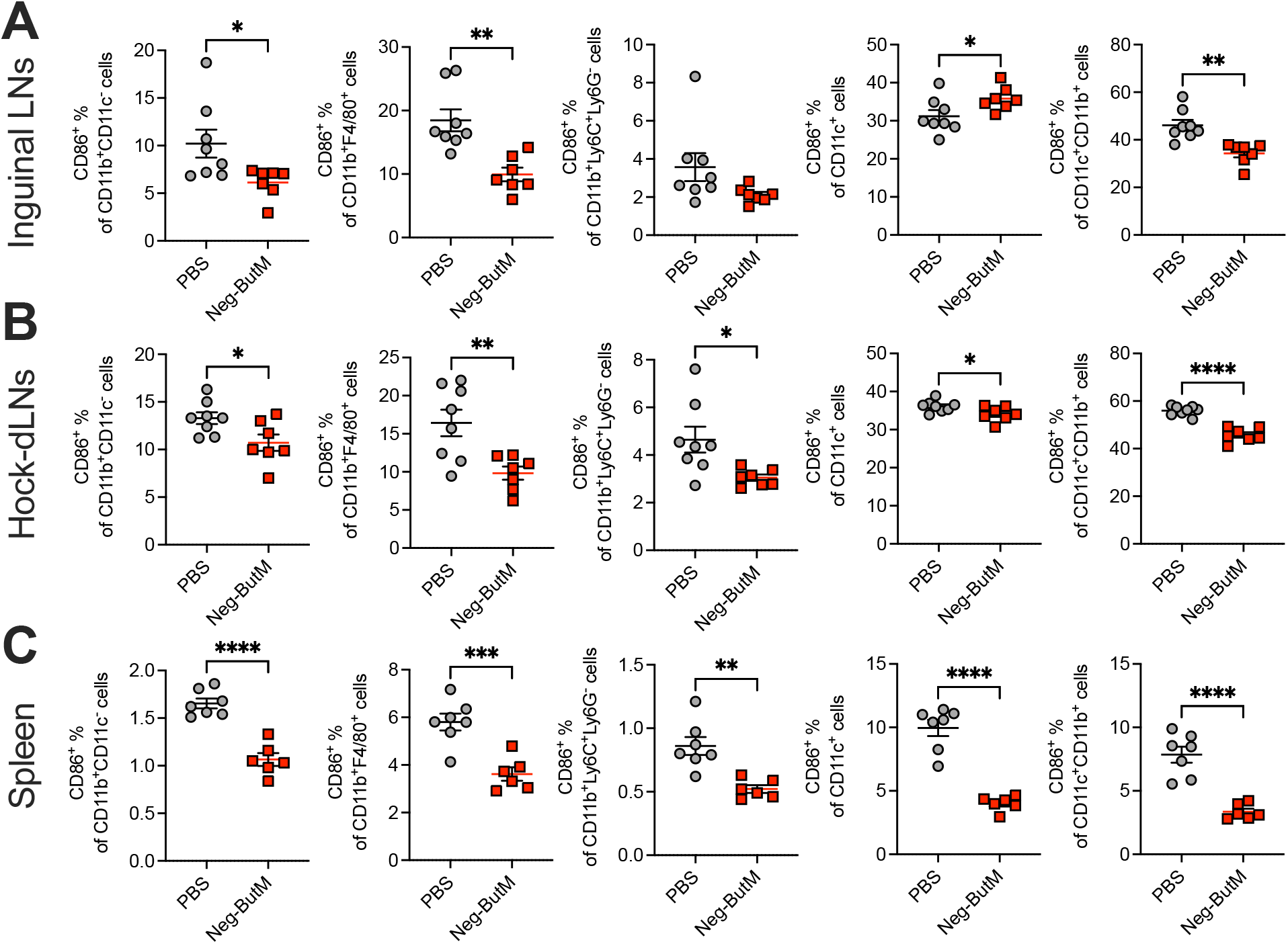
Negatively charged butyrate micelles suppress myeloid cells in arthritis. The percentage of co-stimulatory molecule CD86^+^ of various myeloid cells in the inguinal lymph nodes (LNs) (**A**), hock-draining LNs (**B**), or spleen (**C**), from mice treated with PBS or Neg-ButM in Fig. 4. Data are presented as mean ± SEM. Statistical analyses were performed using Student’s t test. ***p<0.001, ****p<0.0001.

These immune cell characterizations in distinct draining LNs and the spleen demonstrated highly similar outcomes across different s.c. injection sites. We observed concurrent Treg induction, upregulation of PD-1 and CTLA-4 on T cells, and downregulation of CD86 on myeloid cells. These changes could synergistically contribute to a profound suppression of immune activation in the context of CAIA, reinforcing the therapeutic potential of Neg-ButM.

## Discussion

Our previous work developed butyrate prodrug micelles that delivered butyrate to modulate gut health, which included restoring gut barrier permeability and immune homeostasis, following twice daily oral administration in mice(*11*). In this study, we explored these micelles with a focus on directly modulating immune responses through LN targeting after parenteral administration. The findings underscored the significant role of surface charge in influencing LN targeting and cellular uptake, particularly emphasizing the efficacy of negatively charged particles. Following s.c. injection, the negatively charged micelles (Neg-ButM) demonstrated enhanced accumulation and retention not only in LNs but also in other major organs. This facilitated the mitigation of disease progression in collagen antibody-induced arthritis and allowed for the modulation of T cell and myeloid cell responses, both locally and systemically.

Our *in vitro* studies showed that Neg-ButM was more effective at inducing Tregs than Ntl-ButM. This superior efficacy may be attributed to their different rates of uptake: Neg-ButM was taken up more readily across all cell types, especially in CD4^+^ T cells and DCs, which are vital to direct or indirect Treg induction. This finding aligns with previous research suggesting that antigen-presenting cells (APCs), particularly DCs, preferentially engulf negatively charged particles (*48*). Such propensity might mirror natural defense mechanisms against bacteria, which are also negatively charged (*48*). Interestingly, the cellular entry pathways for neutral and negatively charged particles differ. Neutral micelles are generally taken up by clathrin-dependent endocytosis, whereas negative nanoparticles can enter via caveolae-independent pathways (*49*). This distinction may contribute to Neg-ButM’s higher propensity for cellular uptake, observed both *in vitro* and *in vivo* after s.c. administration.

In *in vivo* studies, we showed that Neg-ButM had superior LN accumulation. Nanoparticles have been widely utilized for LN targeting due to their optimal size characteristics for lymphatic drainage (*50*). However, we demonstrated in this study that the charge of these particles also significantly impacts LN targeting and retention. We hypothesized that the anionic microenvironment at the injection site, within the interstitium, drives the preferential drainage of negatively charged nanoparticles into the collecting lymphatic vasculature. In contrast, neutral or positively charged nanoparticles may be trapped within the interstitium. An alternative explanation could be that APCs, especially macrophages, take up more negatively charged nanoparticles, as evidenced by our *in vivo* cellular uptake study, and then transport them into the lymphatic system. The exact mechanism driving this phenomenon warrants further investigation.

Moreover, our study revealed significant immunomodulatory effects of Neg-ButM on both T cells and myeloid cells. Notably, we observed concurrent Treg induction, increased expression of PD-1 and CTLA-4 on both CD4^+^ and CD8^+^ T cells, and downregulation of co-stimulatory markers such as CD86 on macrophages and DCs, both locally and systemically in disease settings. These effects could synergistically contribute to the induction of a tolerogenic environment. However, it remains unclear which cell type plays a more substantial role in Neg-ButM’s potential therapeutic effects. Future studies could use transgenic mouse models to selectively ablate or activate specific immune cell types, or antibodies to deplete specific cell types, to further elucidate this. For example, CD25-neutralizing antibodies could be injected to mice with collagen-induced arthritis to decipher the role of Tregs (*51*, *52*). Clodronate liposomes or anti-CSF1R antibody could be used to deplete macrophages (*53*, *54*), providing insights into their role in mediating the therapeutic effects of Neg-ButM. Utilizing these models could deepen our understanding of Neg-ButM’s therapeutic mechanisms.

In summary, we demonstrated the potential of Neg-ButM in suppressing arthritis, leveraging its direct immunomodulatory effects on the lymphatic system. With just 2-3 s.c. injections, one week apart, Neg-ButM provided long-lasting immunomodulatory effects. Future studies might explore a further decrease in dosage frequency and examine its effects. Although the CAIA model served as an initial proof-of-concept for delivering butyrate to the lymphatic system to suppress autoimmune arthritis prophylactically, this model does not fully recapitulate the human disease pathology, lacking pathogenic T cell or B cell component. While it is challenging to develop mouse models that fully recapitulate human RA, due to the unknown disease-causing antigens, our study nevertheless highlights the promise of developing more effective nano-based treatments for diseases like RA, paving the way for potential applications in treating other autoimmune or chronic inflammatory diseases using long-acting immunomodulation with less frequent dosing regimens.

## Materials and Methods

### Study design

The objective of this study was to explore the impact of surface charge on butyrate micelles’ ability to target LNs, deliver butyrate via a long-acting formulation, induce regulatory T cells, suppress myeloid cells, and test its efficacy in suppressing autoimmune arthritis in mice using the CAIA model.

In our study design, we employed *in vitro* Treg induction models and BMDC stimulation models to establish an initial understanding of the immunomodulatory potential of our micelles. To investigate the biodistribution of our micelles, we compared the LN targeting and cellular uptake capabilities of neutral butyrate micelles (NtL-ButM) and negatively charged butyrate micelles (Neg-ButM) in mice. Antibiotic-treated mice or s.c. LPS-stimulated mice were used to investigate the *in vivo* potential for Treg induction and myeloid cell suppression, respectively. Due to its superior LN targeting and *in vivo* biological effects, Neg-ButM was selected for further efficacy testing in the CAIA model.

In the CAIA model, mice were treated either with a PBS control or Neg-ButM, administered twice with a one-week interval prior to immunization. Paw inflammation was assessed daily, and the pathology of inflamed paws and joints was evaluated at end point using histological techniques. Immune cell population responses were analyzed by flow cytometry following the sacrifice of the mice at the end of the experiment when consistent RA scores had been established.

Sample size determination was based on results obtained from previous and preliminary studies (*55*). In animal studies, each analyzed group consisted of a minimum of 5 and typically 7-8 independent biological replicates. Further details on the sample size for each display figure are shown in the figure legends. All experiments were replicated at least twice for validation. Random assignment was applied to allocate mice to treatment groups. To minimize bias, the person assessing clinical scores for autoimmune arthritis was separate from the person administering treatment and was blinded to the treatment group allocations. Likewise, the person conducting histological analysis was also blinded to treatment groups. Further details on statistical methods can be found in the “Statistical analysis” section.

### Synthesis of NtL-ButM and Neg-ButM

Our group has previously reported the synthesis of two butyrate prodrug micelles bearing different surface charges (*11*). Briefly, Neg-ButM consists of a block copolymer poly(methacrylic acid)-b-poly(butyl methacrylate) (pMAA-b-pBMA). pMAA-b-pBMA was synthesized using the Reversible Addition-Fragmentation chain Transfer (RAFT) polymerization method. The process initiated with the polymerization of the pMAA block from methacrylic acid, followed by the addition of butyl methacrylate to the reaction mixture to form the pBMA block. Neg-ButM micelles were prepared by base titration. The pMAA-b-pBMA polymer was added to PBS under vigorous stirring. A sodium hydroxide (NaOH) solution, molar equivalent to the methacrylic acid, was added to the polymer solution in three portions over 2 hr. After the base solution was added, the polymer solution was stirred at room temperature overnight. Subsequently, PBS was added to reach the target concentration, and the solution was filtered through a 0.22 μm filter, then stored at 4°C for further use.

NtL-ButM is composed of a block copolymer poly(N-(2-hydroxypropyl)methacrylamide)-b-poly(butyl methacrylate) (pHPMA-b-pBMA). The pHPMA-b-pBMA was also synthesized using the RAFT polymerization method. It began with the polymerization of the pHPMA block from N-(2-hydroxypropyl)methacrylamide, followed by the addition of butyl methacrylate to the reaction mixture to polymerize and form the pBMA block. NtL-ButM micelles were formulated using the cosolvent evaporation method. The pHPMA-b-pBMA polymer was dissolved in ethanol at 80 mg/mL with stirring. Once the polymer was fully dissolved, an equal volume of PBS was gradually added. The solution was then allowed to evaporate at room temperature for at least 6 hrs until all the ethanol had been removed. Following evaporation, the NtL-ButM solution was filtered through a 0.22 μm filter and stored at 4°C.

### Mice

C57BL/6 mice, aged 8-12 weeks, were purchased from Charles River (strain code: 027, Charles River). BALB/c mice, aged 6-10 weeks were purchased from the Jackson Laboratory (strain code: 000651, JAX). Foxp3^GFP^ reporter mice were originally purchased from the Jackson Laboratory and then bred in house (strain code: 006769, JAX). All the mice were maintained in a specific pathogen-free (SPF) facility at the University of Chicago. Mice were maintained on a 12 hr light/dark cycle at a room temperature of 20-24°C. All protocols used in this study were approved by the Institutional Animal Care and Use Committee of the University of Chicago.

### Flow cytometry and antibodies

Flow cytometry was done using BD LSRFortessa^TM^, and data were analyzed using FlowJo 10.8.0. Antibodies against the following markers were used in this study: CD3 (APC-Cy7, Cat#100222, BioLegend), CD3 (BV605, Cat#100237, BioLegend), CD3 (BUV737, Cat#741788, BD Biosciences), B220 (BV421, Cat#103251, BioLegend), CD4 (FITC Cat#116003, BioLegend), CD4 (BV786, Cat#740844, BD Biosciences), CD4 (BV605, Cat#100548, BioLegend), CD8 (AF700, Cat#100730, BioLegend), F4/80 (PE, Cat#123110, BioLegend), F4/80 (APC-Cy7, Cat#123117, BioLegend), F4/80 (APC, Cat#123116, BioLegend), CD11b (PerCP-Cy5.5, Cat#550993, BD Biosciences), CD11b (BV786, Cat#101243, BioLegend), CD11b (BV711, Cat#101242, BioLegend), CD11c (BV421, Cat#562782, BD Biosciences), CD11c (BV786, Cat#117335, BioLegend), CD11c (PE-Cy7, Cat#558079, BD Biosciences), CD11c (PE, Cat#117308, BioLegend), CD25 (PE, Cat#561065, BD Biosciences), CD25 (APC, Cat#162105, BioLegend), CD40 (PerCP-Cy5.5, Cat#124623, BioLegend), I-A/I-E (FITC, Cat#107606, BioLegend), CD169 (BV605, Cat#142413, BioLegend), Foxp3 (AF488, Cat#53-5773-82, Invitrogen), PD-1 (BV711, Cat#135231, BioLegend), CTLA-4 (PE-Cy7, Cat#25-1522-80, Invitrogen), Ly6C (BV605, Cat#128036, BioLegend), Ly6G (AF488, Cat#127626, BioLegend), CD86 (AF700, Cat#105024, BioLegend).

### *In vitro* Treg induction

Splenocytes were isolated from 5-7 wk old male or female Foxp3^GFP^ reporter mice, and purified using lysing buffer. The isolated cells were then seeded at 500,000 cells per well in a round-bottom 96 well plate in IMDM medium (Life Technologies), supplemented with 10% HIFBS (Gibco) and 1% penicillin/streptomycin (Sigma-Aldrich). To create an environment conducive to Treg induction and expansion, IL-2 (10 ng/mL), TGFβ (0.2 ng/mL), anti-CD3ε (10 µg/mL) and anti-CD28 (1 µg/mL) were added to all wells. Furthermore, either NtL-ButM or Neg-ButM were added at concentrations equivalent to butyrate of 0.5, 1, 2, or 4 mM. The cells were incubated for 4 days, after which they were collected, stained with live/dead stain (Cat#L34957, Invitrogen) and fluorescent antibodies against CD4 and CD25. Finally, the cells were analyzed using flow cytometry (BD LSRFortessa).

### Mouse BMDC isolation and activation study

Murine BMDCs were obtained from 6-week-old female C57BL/6 mice as previously described by Lutz et al (*56*). The BMDCs were seeded at a density of 3 × 10^6^ cells/plate in petri dishes, and cultured at 37°C and 5% CO_2_ in RPMI 1640 medium (Life Technologies) supplemented with 10% HIFBS (Gibco), GM-CSF (20 ng/mL; recombinant Mouse GM-CSF (carrier-free) from BioLegend), 2 mM l-glutamine (Life Technologies), 1% antibiotic-antimycotic (Life Technologies). The culture medium was replenished on day 3 and day 6, and cells were used on day 9. Isolated BMDCs were then seeded in round-bottom 96 well plates at 100,000 cells per well in RPMI media. The BMDCs were incubated with varing concentrations of either NtL-ButM or Neg-ButM at concentrations equivalent to butyrate of 0.5, 1, 2, or 4 mM for 24 hrs. After this period, LPS (1 μg/mL) was added for an additional18 hr. The supernatant of cell culture was collected for cytokine analysis using LEGENDPlex^TM^ (BioLegend). The BMDCs were also collected and stained with live/dead stain (Cat#L34957, Invitrogen) and fluorescent antibodies against CD11c, MHCII, and CD40, and subsequently analyzed via flow cytometry (BD LSRFortessa).

### Biodistribution of NtL-ButM and Neg-ButM

SPF C57BL/6 mice were used for *in vivo* biodistribution studies. Azide-labelled pHPMA-b-pBMA or pMAA-b-pBMA polymer was reacted with IR 750-DBCO (Fluoroprobes) and purified via hexane precipitation. Following formulation into micelles, as described previously, the fluorescently labeled NtL-ButM or Neg-ButM were s.c. administered to mice at the right abdominal site. At 7 d or 35 d post-administration, mice were euthanized, and their right and left inguinal LNs, small intestine or colon-draining LNs, as well as the major organs were harvested. The whole-organ fluorescence was measured via an In Vivo Imaging System (IVIS) (Perkin Elmer). Images were processed and analyzed using Living Imaging 4.5.5 (Perkin Elmer).

In the cellular biodistribution study, Azide-labelled pHPMA-b-pBMA or pMAA-b-pBMA polymer was reacted with AFDye647-DBCO (Click Chemistry Tools) and then formulated into micelles. The fluorescently labeled NtL-ButM or Neg-ButM were administered s.c. to mice in the right abdominal site. Mice were euthanized at 1 d or 3 d post-administration. Right and left inguinal LNs, small intestine-draining LNs, and spleen were harvested. Cells were isolated from these tissues into single cell suspensions, stained with live/dead and fluorescent antibodies against B220, CD3, CD4, F4/80, CD11b, and CD11c, and analyzed by the flow cytometry.

### Treg induction and myeloid cell suppression study on healthy mice

For *in vivo* Treg induction study, starting at weaning, Foxp3GFP reporter mice were given drinking water containing a mixture of antibiotics, including ampicillin (1 mg/mL), neomycin (1 mg/mL), metronidazole (0.2 mg/mL), and vancomycin (0.5 mg/mL) (Sigma-Aldrich) starting at weaning. This antibiotic cocktail composition and treatment duration are consistent with standard protocols for depleting microbiota, thus eliminating the influence of endogenous butyrate (*57–59*). On day 4 post-weaning, mice were s.c. administered in the abdominal site with saline, NaBut, NtL-ButM, or Neg-ButM at the equivalent butyrate dose of 2.24 mg per mice once a week for three weeks. One week after the final dose, mice were sacrificed and cells from the draining inguinal LNs were isolated and stained with live/dead and fluorescent antibodies against CD3, CD4, and CD25, followed by the flow cytometry analysis.

For the *in vivo* myeloid cell suppression study, C57BL/6 mice were s.c. administered at the abdominal site with saline, NaBut, NtL-ButM, or Neg-ButM at the equivalent butyrate dose of 2.24 mg per mouse on day 0. On day 6, mice were injected with LPS s.c. at the same injection site. On day 7, mice were sacrificed and cells from the draining inguinal LNs were isolated and stained with live/dead and fluorescent antibodies against CD11c, I-A/I-E, CD169, CD11b, and F4/80, followed by flow cytometry analysis.

### Collagen-antibody inducing arthritis (CAIA) model

BALB/c mice at age of 8-10 weeks were s.c. injected with either PBS or Neg-ButM (equivalent to 2.8 mg butyrate per mice) on day -8 and day -1. CAIA was induced by passive immunization with an anti-collagen antibody cocktail (1 mg per mice, Arthrogen-CIA® 5-Clone Cocktail Kit, Chondrex, Inc.) on day 0, followed by an intraperitoneal injection of LPS (25 μg) on day 3. Mice were provided with soft (pine) bedding throughout the experiment. Arthritis severity was monitored daily after day 3 according to the criteria for clinical scores as instructed by Chondrex, Inc., as described previously (*55*, *60*).

On day 12, the thickness of mouse fore and hindpaws was measured to assess the swelling resulting from arthritis. On day 13, mice were sacrificed, and the spleen, inguinal LNs, and hock-draining LNs, including popliteal, axillary, and brachial LNs, were harvested for immunostaining and subsequent flow cytometry analysis. Paws were collected for histological analysis, as previously described (*55*). Briefly, paws were fixed in 2% paraformaldehyde (Thermo Scientific), decalcified in Decalcifer II (Leica), and stored in 70% ethanol until paraffin embedding. Paraffin-embedded paws were sliced into 5 μm-thick sections and stained with hematoxylin and eosin (H&E). Images were scanned using an Olympus VS200 Slideview Research Slide Scanner and analyzed using QuPath software.

### Statistical analysis

Statistical analysis and plotting of data were performed using Prism 10.0 (Graphpad), as indicated in the figure legends. One-way ANOVA with Dunnett’s or Kruskal-Wallis test (if not normally distributed) for multiple comparisons was used in **Fig. 1E, H, I-K, and Fig. 3B, D**. Student’s t-test was used in **Fig. 4B-E**, **Fig. 5, and Fig. 6**. Two-way ANOVA with Bonferroni’s post-test was used in **Fig. 1C, F**, **Fig. 2C, D**. In **Fig. 4B**, the area under curve (AUC) values of clinical scores were compared using Student’s t test. Data represent mean ± SEM; *n* is stated in the figure legend. *p<0.05, **p<0.01, ***p<0.001, ****p<0.0001.

## Acknowledgements

This work was supported by the Chicago Immunoengineering Innovation Center of the University of Chicago and the Alper Family Fund. We thank Prof. Cathryn R. Nagler at the University of Chicago for providing Foxp3^GFP^ reporter mice, as well as her and other Nagler lab members for providing constructive feedback on this project. We thank Yue Wang from Prof. Melody A. Swartz’s laboratory for providing mouse BMDCs. We thank Suzana Gomes for the endotoxin testing on all the materials that we injected to mice. We thank the Cytometry and Antibody Technology Core Facility (Cancer Center Support Grant P30CA014599), and the Human Tissue Resource Center, the Integrated Light Microscopy Core at the University of Chicago.

## Author contributions

J.A.H. oversaw all research. S.C., E.B., and J.A.H. designed most of experiments. S.C., and R. W. synthesized materials. S.C., E.B., R.W., M.S., A.S., M.N., K.H., A.D. performed experiments. S.C. and E.B. analyzed experiments. S.C., and J.A.H. wrote the manuscript. All authors contributed to the article and approved the submitted version.

## References

1. T. M. Hunter, N. N. Boytsov, X. Zhang, K. Schroeder, K. Michaud, A. B. Araujo, Prevalence of rheumatoid arthritis in the United States adult population in healthcare claims databases, 2004-2014. Rheumatol. Int. 37, 1551–1557 (2017).

2. S. Shams, J. M. Martinez, J. R. D. Dawson, J. Flores, M. Gabriel, G. Garcia, A. Guevara, K. Murray, N. Pacifici, M. V. Vargas, T. Voelker, J. W. Hell, J. F. Ashouri, The Therapeutic Landscape of Rheumatoid Arthritis: Current State and Future Directions. Front. Pharmacol. 12 (2021).

3. V. M. Holers, M. K. Demoruelle, K. A. Kuhn, J. H. Buckner, W. H. Robinson, Y. Okamoto, J. M. Norris, K. D. Deane, Rheumatoid arthritis and the mucosal origins hypothesis: protection turns to destruction. Nat. Rev. Rheumatol. 14, 542–557 (2018).

4. A. I. Catrina, K. D. Deane, J. U. Scher, Gene, environment, microbiome and mucosal immune tolerance in rheumatoid arthritis. Rheumatology. 55, 391–402 (2016).

5. Y. Maeda, K. Takeda, Role of Gut Microbiota in Rheumatoid Arthritis. J. Clin. Med. 6, 60 (2017).

6. M. M. Zaiss, H.-J. Joyce Wu, D. Mauro, G. Schett, F. Ciccia, The gut–joint axis in rheumatoid arthritis. Nat. Rev. Rheumatol. 17, 224–237 (2021).

7. G. Horta-Baas, M. D. S. Romero-Figueroa, A. J. Montiel-Jarquín, M. L. Pizano-Zárate, J. García-Mena, N. Ramírez-Durán, Intestinal Dysbiosis and Rheumatoid Arthritis: A Link between Gut Microbiota and the Pathogenesis of Rheumatoid Arthritis. J. Immunol. Res. 2017, 4835189 (2017).

8. E. C. Rosser, C. J. M. Piper, D. E. Matei, P. A. Blair, A. F. Rendeiro, M. Orford, D. G. Alber, T. Krausgruber, D. Catalan, N. Klein, J. J. Manson, I. Drozdov, C. Bock, L. R. Wedderburn, S. Eaton, C. Mauri, Microbiota-Derived Metabolites Suppress Arthritis by Amplifying Aryl-Hydrocarbon Receptor Activation in Regulatory B Cells. Cell Metab. 31, 837–851.e10 (2020).

9. D. R. Donohoe, N. Garge, X. Zhang, W. Sun, T. M. O’Connell, M. K. Bunger, S. J. Bultman, The Microbiome and Butyrate Regulate Energy Metabolism and Autophagy in the Mammalian Colon. Cell Metab. 13, 517–526 (2011).

10. J. M. Wells, R. J. Brummer, M. Derrien, T. T. MacDonald, F. Troost, P. D. Cani, V. Theodorou, J. Dekker, A. Méheust, W. M. de Vos, A. Mercenier, A. Nauta, C. L. Garcia-Rodenas, Homeostasis of the gut barrier and potential biomarkers. Am. J. Physiol.-Gastrointest. Liver Physiol. 312, G171–G193 (2017).

11. R. Wang, S. Cao, M. E. H. Bashir, L. A. Hesser, Y. Su, S. M. C. Hong, A. Thompson, E. Culleen, M. Sabados, N. P. Dylla, E. Campbell, R. Bao, E. B. Nonnecke, C. L. Bevins, D. S. Wilson, J. A. Hubbell, C. R. Nagler, Treatment of peanut allergy and colitis in mice via the intestinal release of butyrate from polymeric micelles. *Nat*. Biomed. Eng., 1–18 (2022).

12. A. Koh, F. De Vadder, P. Kovatcheva-Datchary, F. Bäckhed, From Dietary Fiber to Host Physiology: Short-Chain Fatty Acids as Key Bacterial Metabolites. Cell. 165, 1332–1345 (2016).

13. J. A. McKay, J. C. Mathers, Diet induced epigenetic changes and their implications for health. Acta Physiol. Oxf. Engl. 202, 103–118 (2011).

14. R. Berni Canani, M. Di Costanzo, L. Leone, The epigenetic effects of butyrate: potential therapeutic implications for clinical practice. Clin. Epigenetics. 4, 4 (2012).

15. J. Tan, C. McKenzie, M. Potamitis, A. N. Thorburn, C. R. Mackay, L. Macia, The role of short-chain fatty acids in health and disease. Adv. Immunol. 121, 91–119 (2014).

16. H. M. Hamer, D. Jonkers, K. Venema, S. Vanhoutvin, F. J. Troost, R.-J. Brummer, Review article: the role of butyrate on colonic function. Aliment. Pharmacol. Ther. 27, 104–119 (2008).

17. P. M. Smith, M. R. Howitt, N. Panikov, M. Michaud, C. A. Gallini, M. Bohlooly-Y, J. N. Glickman, W. S. Garrett, The microbial metabolites, short-chain fatty acids, regulate colonic Treg cell homeostasis. Science. 341, 569–573 (2013).

18. Y. Furusawa, Y. Obata, S. Fukuda, T. A. Endo, G. Nakato, D. Takahashi, Y. Nakanishi, C. Uetake, K. Kato, T. Kato, M. Takahashi, N. N. Fukuda, S. Murakami, E. Miyauchi, S. Hino, K. Atarashi, S. Onawa, Y. Fujimura, T. Lockett, J. M. Clarke, D. L. Topping, M. Tomita, S. Hori, O. Ohara, T. Morita, H. Koseki, J. Kikuchi, K. Honda, K. Hase, H. Ohno, Commensal microbe-derived butyrate induces the differentiation of colonic regulatory T cells. Nature. 504, 446–450 (2013).

19. N. Arpaia, C. Campbell, X. Fan, S. Dikiy, J. van der Veeken, P. deRoos, H. Liu, J. R. Cross, K. Pfeffer, P. J. Coffer, A. Y. Rudensky, Metabolites produced by commensal bacteria promote peripheral regulatory T-cell generation. Nature. 504, 451–455 (2013).

20. M. M. M. Kaisar, L. R. Pelgrom, A. J. van der Ham, M. Yazdanbakhsh, B. Everts, Butyrate Conditions Human Dendritic Cells to Prime Type 1 Regulatory T Cells via both Histone Deacetylase Inhibition and G Protein-Coupled Receptor 109A Signaling. Front. Immunol. 8, 1429 (2017).

21. C. Nastasi, M. Candela, C. M. Bonefeld, C. Geisler, M. Hansen, T. Krejsgaard, E. Biagi, M. H. Andersen, P. Brigidi, N. Ødum, T. Litman, A. Woetmann, The effect of short-chain fatty acids on human monocyte-derived dendritic cells. Sci. Rep. 5, 16148 (2015).

22. Z. Ang, J. L. Ding, GPR41 and GPR43 in Obesity and Inflammation – Protective or Causative? Front. Immunol. 7 (2016).

23. A. J. Brown, S. M. Goldsworthy, A. A. Barnes, M. M. Eilert, L. Tcheang, D. Daniels, A. I. Muir, M. J. Wigglesworth, I. Kinghorn, N. J. Fraser, N. B. Pike, J. C. Strum, K. M. Steplewski, P. R. Murdock, J. C. Holder, F. H. Marshall, P. G. Szekeres, S. Wilson, D. M. Ignar, S. M. Foord, A. Wise, S. J. Dowell, The Orphan G protein-coupled receptors GPR41 and GPR43 are activated by propionate and other short chain carboxylic acids. J. Biol. Chem. 278, 11312–11319 (2003).

24. R. I. Breuer, K. H. Soergel, B. A. Lashner, M. L. Christ, S. B. Hanauer, A. Vanagunas, J. M. Harig, A. Keshavarzian, M. Robinson, J. H. Sellin, D. Weinberg, D. E. Vidican, K. L. Flemal, A. W. Rademaker, Short chain fatty acid rectal irrigation for left-sided ulcerative colitis: a randomised, placebo controlled trial. Gut. 40, 485–491 (1997).

25. G. D. Sher, G. D. Ginder, J. Little, S. Yang, G. J. Dover, N. F. Olivieri, Extended therapy with intravenous arginine butyrate in patients with beta-hemoglobinopathies. N. Engl. J. Med. 332, 1606–1610 (1995).

26. P. Vernia, M. Cittadini, R. Caprilli, A. Torsoli, Topical treatment of refractory distal ulcerative colitis with 5-ASA and sodium butyrate. Dig. Dis. Sci. 40, 305–307 (1995).

27. H. Liu, J. Wang, T. He, S. Becker, G. Zhang, D. Li, X. Ma, Butyrate: A Double-Edged Sword for Health? Adv. Nutr. Bethesda Md. 9, 21–29 (2018).

28. N. L. Trevaskis, L. M. Kaminskas, C. J. H. Porter, From sewer to saviour — targeting the lymphatic system to promote drug exposure and activity. Nat. Rev. Drug Discov. 14, 781– 803 (2015).

29. W. Xu, N. R. Harris, K. M. Caron, Lymphatic Vasculature: An Emerging Therapeutic Target and Drug Delivery Route. Annu. Rev. Med. 72, 167–182 (2021).

30. J. McCright, R. Naiknavare, J. Yarmovsky, K. Maisel, Targeting Lymphatics for Nanoparticle Drug Delivery. Front. Pharmacol. 13, 887402 (2022).

31. S. Cao, S. D. Slack, C. N. Levy, S. M. Hughes, Y. Jiang, C. Yogodzinski, P. Roychoudhury, K. R. Jerome, J. T. Schiffer, F. Hladik, K. A. Woodrow, Hybrid nanocarriers incorporating mechanistically distinct drugs for lymphatic CD4+ T cell activation and HIV-1 latency reversal. Sci. Adv. 5, eaav6322 (2019).

32. H. Jiang, Q. Wang, X. Sun, Lymph node targeting strategies to improve vaccination efficacy. J. Controlled Release. 267, 47–56 (2017).

33. M. Liu, S. Li, M. O. Li, TGF-β Control of Adaptive Immune Tolerance: A Break From Treg Cells. BioEssays. 40, 1800063 (2018).

34. J. Zhao, J. Zhao, S. Perlman, Differential effects of IL-12 on Tregs and non-Treg T cells: roles of IFN-γ, IL-2 and IL-2R. PloS One. 7, e46241 (2012).

35. F. S. Kleijwegt, S. Laban, G. Duinkerken, A. M. Joosten, A. Zaldumbide, T. Nikolic, B. O. Roep, Critical role for TNF in the induction of human antigen-specific regulatory T cells by tolerogenic dendritic cells. J. Immunol. Baltim. Md 1950. 185, 1412–1418 (2010).

36. C. Barthels, A. Ogrinc, V. Steyer, S. Meier, F. Simon, M. Wimmer, A. Blutke, T. Straub, U. Zimber-Strobl, E. Lutgens, P. Marconi, C. Ohnmacht, D. Garzetti, B. Stecher, T. Brocker, CD40-signalling abrogates induction of RORγt+ Treg cells by intestinal CD103+ DCs and causes fatal colitis. Nat. Commun. 8, 14715 (2017).

37. T. Tanaka, M. Narazaki, T. Kishimoto, IL-6 in Inflammation, Immunity, and Disease. Cold Spring Harb. Perspect. Biol. 6, a016295 (2014).

38. H. Wiig, M. A. Swartz, Interstitial Fluid and Lymph Formation and Transport: Physiological Regulation and Roles in Inflammation and Cancer. Physiol. Rev. 92, 1005–1060 (2012).

39. S. Cao, Y. Jiang, C. N. Levy, S. M. Hughes, H. Zhang, F. Hladik, K. A. Woodrow, Optimization and comparison of CD4-targeting lipid–polymer hybrid nanoparticles using different binding ligands. J. Biomed. Mater. Res. A. 106, 1177–1188 (2018).

40. M. L. Stoll, R. Kumar, C. D. Morrow, E. J. Lefkowitz, X. Cui, A. Genin, R. Q. Cron, C. O. Elson, Altered microbiota associated with abnormal humoral immune responses to commensal organisms in enthesitis-related arthritis. Arthritis Res. Ther. 16, 486 (2014).

41. D. S. Kim, J.-E. Kwon, S. H. Lee, E. K. Kim, J.-G. Ryu, K.-A. Jung, J.-W. Choi, M.-J. Park, Y.-M. Moon, S.-H. Park, M.-L. Cho, S.-K. Kwok, Attenuation of Rheumatoid Inflammation by Sodium Butyrate Through Reciprocal Targeting of HDAC2 in Osteoclasts and HDAC8 in T Cells. Front. Immunol. 9, 1525 (2018).

42. K. Terato, D. S. Harper, M. M. Griffiths, D. L. Hasty, X. J. Ye, M. A. Cremer, J. M. Seyer, Collagen-induced arthritis in mice: synergistic effect of E. coli lipopolysaccharide bypasses epitope specificity in the induction of arthritis with monoclonal antibodies to type II collagen. Autoimmunity. 22, 137–147 (1995).

43. U. Schulze-Topphoff, M. Varrin-Doyer, K. Pekarek, C. M. Spencer, A. Shetty, S. A. Sagan, B. A. C. Cree, R. A. Sobel, B. T. Wipke, L. Steinman, R. H. Scannevin, S. S. Zamvil, Dimethyl fumarate treatment induces adaptive and innate immune modulation independent of Nrf2. Proc. Natl. Acad. Sci. U. S. A. 113, 4777–4782 (2016).

44. B. Zhu, Y. Bando, S. Xiao, K. Yang, A. C. Anderson, V. K. Kuchroo, S. J. Khoury, CD11b+Ly-6C(hi) suppressive monocytes in experimental autoimmune encephalomyelitis. J. Immunol. Baltim. Md 1950. 179, 5228–5237 (2007).

45. D. M. Sansom, CD28, CTLA-4 and their ligands: who does what and to whom? Immunology. 101, 169–177 (2000).

46. P. S. Linsley, J. A. Ledbetter, The role of the CD28 receptor during T cell responses to antigen. Annu. Rev. Immunol. 11, 191–212 (1993).

47. L. S. K. Walker, D. M. Sansom, The emerging role of CTLA4 as a cell-extrinsic regulator of T cell responses. Nat. Rev. Immunol. 11, 852–863 (2011).

48. E. Fröhlich, The role of surface charge in cellular uptake and cytotoxicity of medical nanoparticles. Int. J. Nanomedicine. 7, 5577–5591 (2012).

49. G. Sahay, D. Y. Alakhova, A. V. Kabanov, Endocytosis of nanomedicines. J. Control. Release Off. J. Control. Release Soc. 145, 182–195 (2010).

50. A. Schudel, D. M. Francis, S. N. Thomas, Material design for lymph node drug delivery. Nat. Rev. Mater. 4, 415–428 (2019).

51. M. E. Morgan, R. P. M. Sutmuller, H. J. Witteveen, L. M. van Duivenvoorde, E. Zanelli, C. J. M. Melief, A. Snijders, R. Offringa, R. R. P. de Vries, R. E. M. Toes, CD25+ cell depletion hastens the onset of severe disease in collagen-induced arthritis. Arthritis Rheum. 48, 1452–1460 (2003).

52. D. S. Wilson, M. Damo, S. Hirosue, M. M. Raczy, K. Brünggel, G. Diaceri, X. Quaglia-Thermes, J. A. Hubbell, Synthetically glycosylated antigens induce antigen-specific tolerance and prevent the onset of diabetes. *Nat*. Biomed. Eng. 3, 817–829 (2019).

53. N. van Rooijen, E. van Kesteren-Hendrikx, Clodronate Liposomes: Perspectives in Research and Therapeutics. J. Liposome Res. 12, 81–94 (2002).

54. C. D. Maulloo, S. Cao, E. A. Watkins, M. M. Raczy, Ani. S. Solanki, M. Nguyen, J. W. Reda, H.-N. Shim, D. S. Wilson, M. A. Swartz, J. A. Hubbell, Lymph Node-Targeted Synthetically Glycosylated Antigen Leads to Antigen-Specific Immunological Tolerance. Front. Immunol. 12 (2021).

55. E. Yuba, E. Budina, K. Katsumata, A. Ishihara, A. Mansurov, A. T. Alpar, E. A. Watkins, P. Hosseinchi, J. W. Reda, A. L. Lauterbach, M. Nguyen, A. Solanki, T. Kageyama, M. A. Swartz, J. Ishihara, J. A. Hubbell, Suppression of Rheumatoid Arthritis by Enhanced Lymph Node Trafficking of Engineered Interleukin-10 in Murine Models. Arthritis Rheumatol. 73, 769–778 (2021).

56. M. B. Lutz, N. Kukutsch, A. L. J. Ogilvie, S. Rößner, F. Koch, N. Romani, G. Schuler, An advanced culture method for generating large quantities of highly pure dendritic cells from mouse bone marrow. J. Immunol. Methods. 223, 77–92 (1999).

57. S. Rakoff-Nahoum, J. Paglino, F. Eslami-Varzaneh, S. Edberg, R. Medzhitov, Recognition of Commensal Microflora by Toll-Like Receptors Is Required for Intestinal Homeostasis. Cell. 118, 229–241 (2004).

58. A. Morgun, A. Dzutsev, X. Dong, R. L. Greer, D. J. Sexton, J. Ravel, M. Schuster, W. Hsiao, P. Matzinger, N. Shulzhenko, Uncovering effects of antibiotics on the host and microbiota using transkingdom gene networks. Gut. 64, 1732–1743 (2015).

59. S. Cao, C. D. Maulloo, M. M. Raczy, M. Sabados, A. J. Slezak, M. Nguyen, A. Solanki, R. P. Wallace, H.-N. Shim, D. S. Wilson, J. A. Hubbell, Glycopolymer-conjugated antigens as an inverse vaccine platform prevent anaphylaxis in a pre-clinical model of food allergy. BioRxiv (2023), p. 2023.03.23.534004, doi:10.1101/2023.03.23.534004.

60. S. Cao, E. Budina, M. M. Raczy, A. Solanki, M. Nguyen, K. Hultgren, P. Ang, J. W. Reda, L. S. Shores, I. Pillai, R. P. Wallace, A. Dhar, E. A. Watkins, J. A. Hubbell, Seryl-butyrate: a prodrug that enhances butyrate’s oral bioavailability and suppresses autoimmune arthritis and experimental autoimmune encephalomyelitis. BioRxiv (2023), p. 2023.04.28.538720, doi:10.1101/2023.04.28.538720.

