## Supplementary Information for "Injectable butyrate-prodrug micelles induce long-acting immune modulation and suppress autoimmune arthritis in mice"

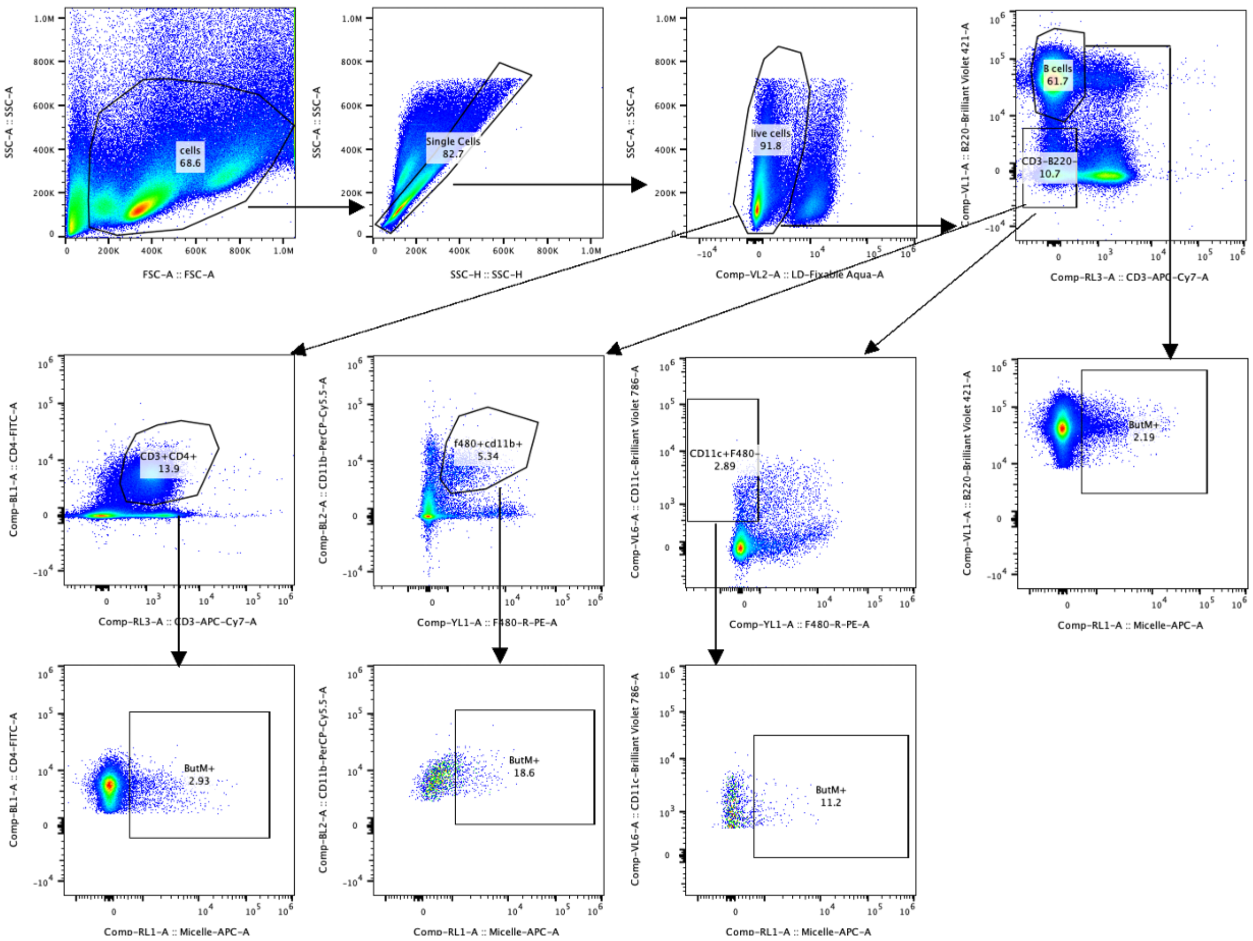


**Figure S1.** Flow cytometry gating strategy for the identification of fluorescently-labeled ButM$^+$ cells of CD4$^+$ T cells, B cells, F4/80$^+$CD11b$^+$ macrophages, and CD11c$^+$ dendritic cells. This gating strategy was used in Fig.1C, F and Fig.2C, D.

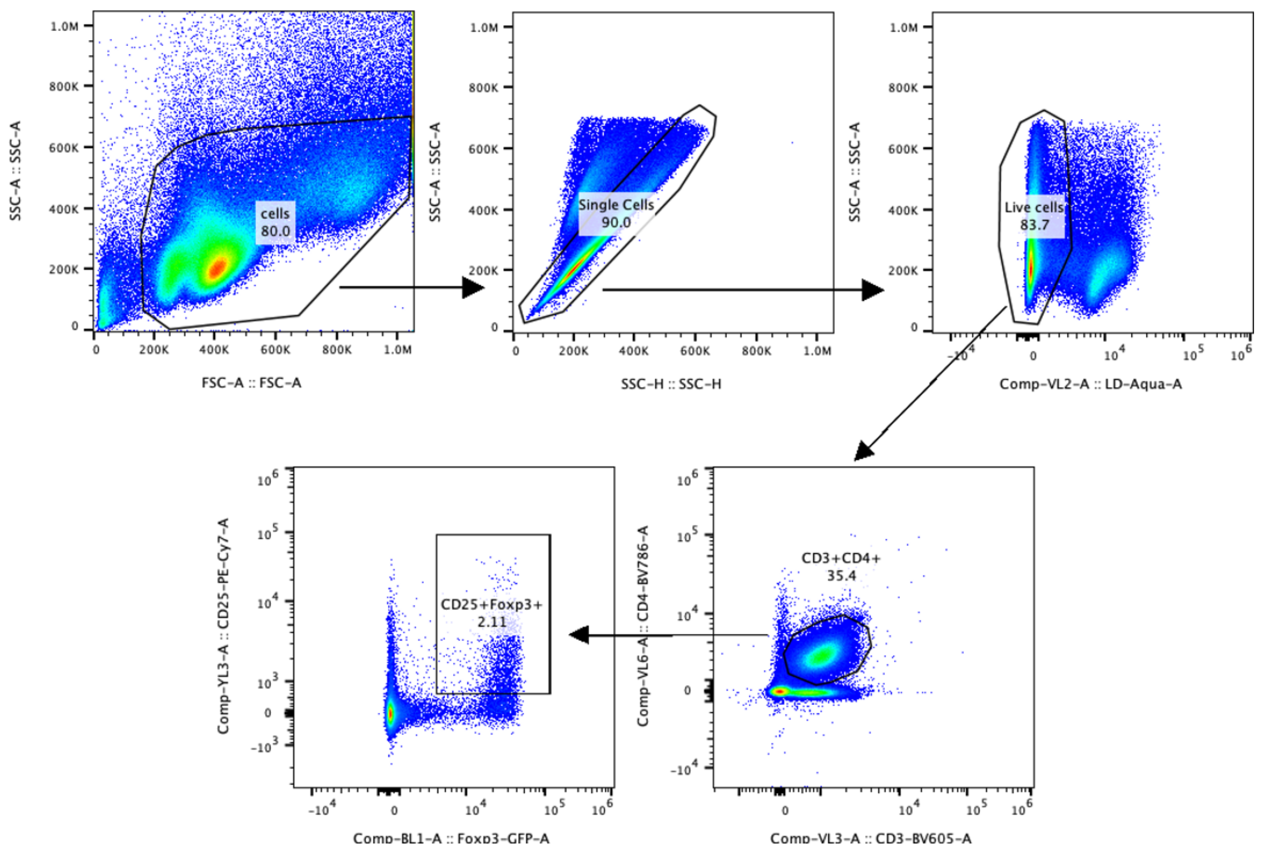


**Figure S2.** Flow cytometry gating strategy for the identification of Foxp3(GFP)$^{+}$CD25$^{+}$ regulatory CD4$^{+}$ T cells. This gating strategy was used in Fig.1E and Fig.3B.

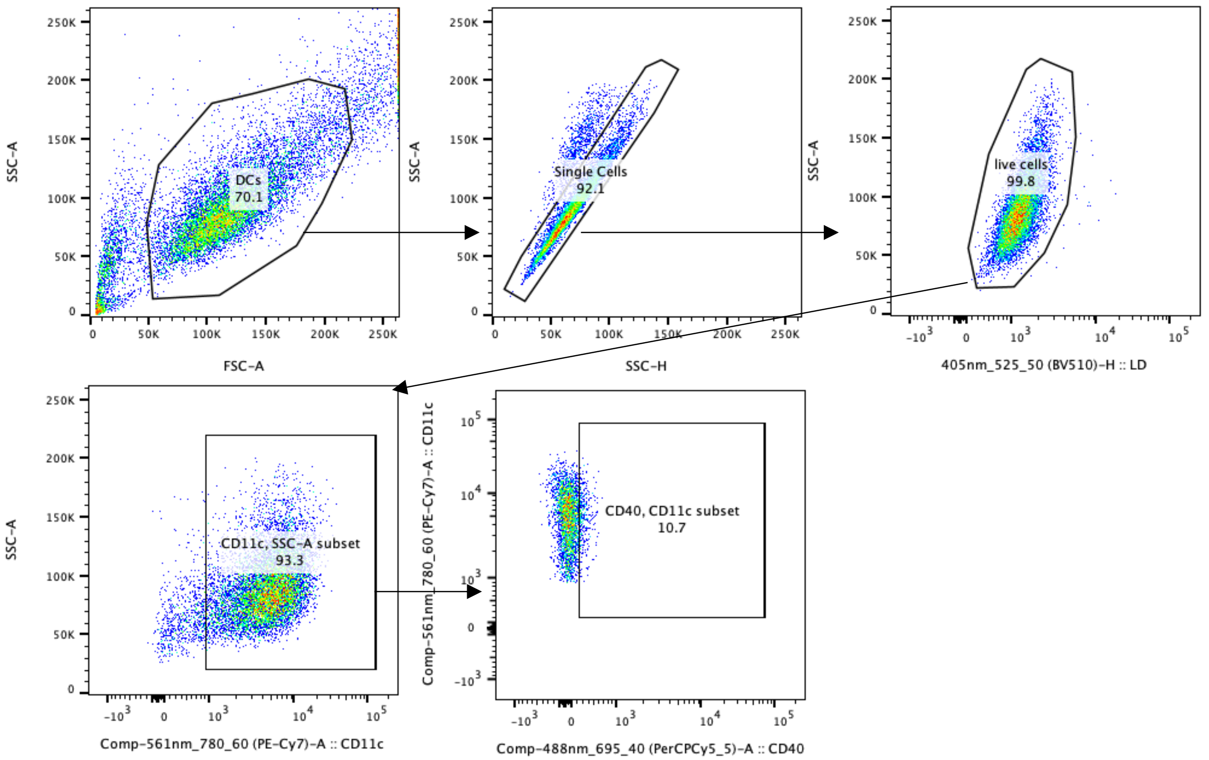


**Figure S3.** Flow cytometry gating strategy for the identification of CD40$^+$ cells of mouse bone marrow-derived dendritic cells (BMDCs). This gating strategy was used in Fig.1H.

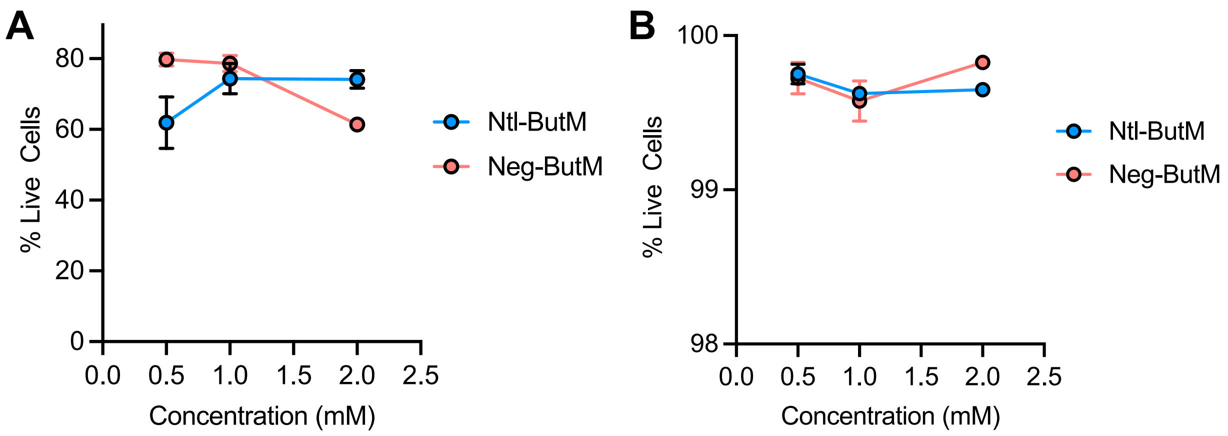


**Figure S4.** Percentage of live cells after treatment with NtL-ButM or Neg-ButM. **A.** Splenocytes were isolated from Foxp3GFP reporter mice and treated with NtL-ButM or Neg-ButM for 4 days. **B.** BMDCs were isolated from C57BL/6 mice and treated with NtL-ButM or Neg-ButM for 24 hr, followed by the addition of LPS for an additional 18 hrs.

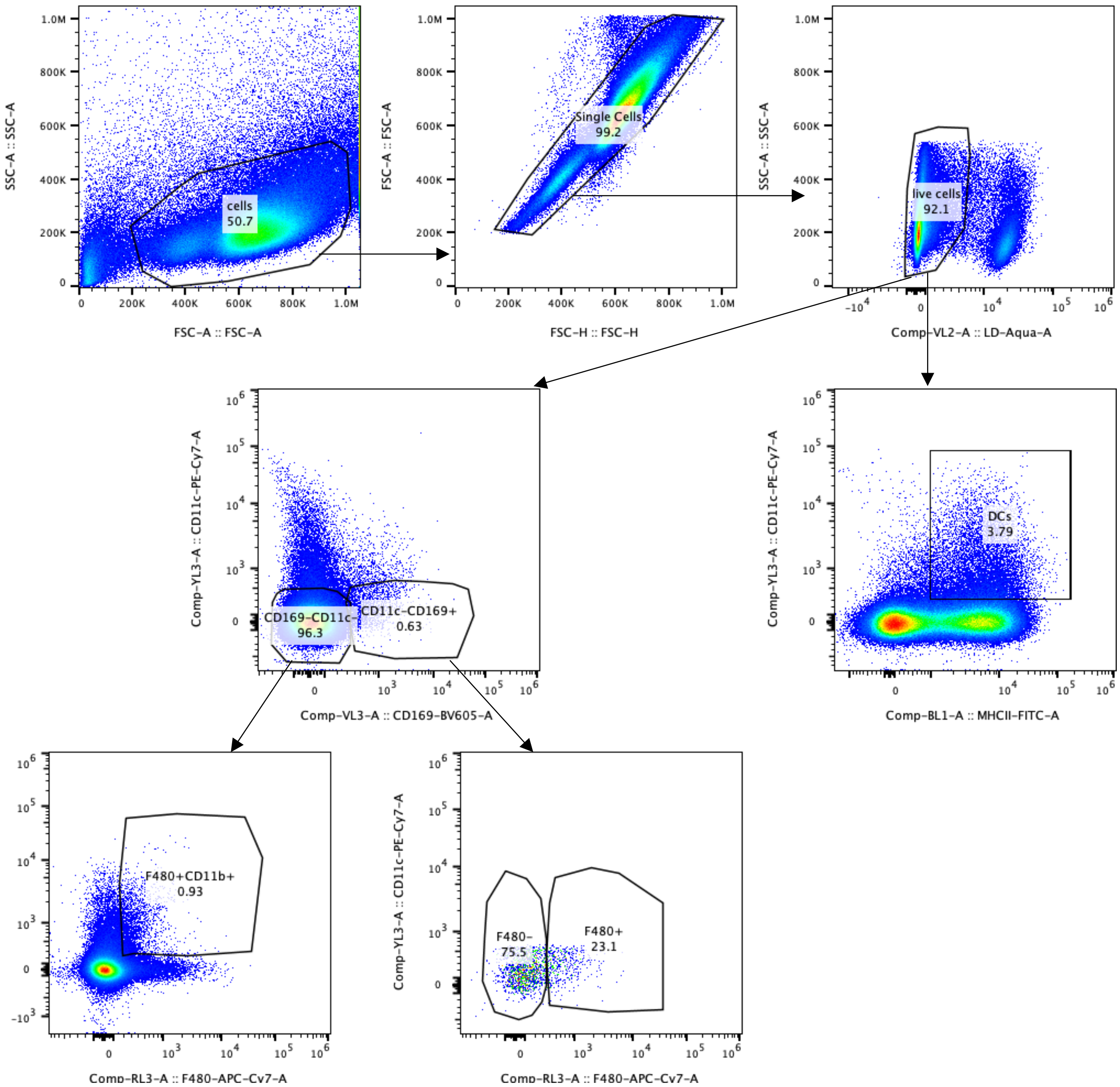


**Figure S5.** Flow cytometry gating strategy for $CD11b^+F4/80^+$ macrophages, subcapsular macrophages ($CD169^+$ $CD11c^{low}F4/80^-$), marginal zone macrophages ($CD169^+$ $CD11c^{low}F4/80^+$), and $CD11b^+$ dendritic cells. This gating strategy was used in Fig. 3D.

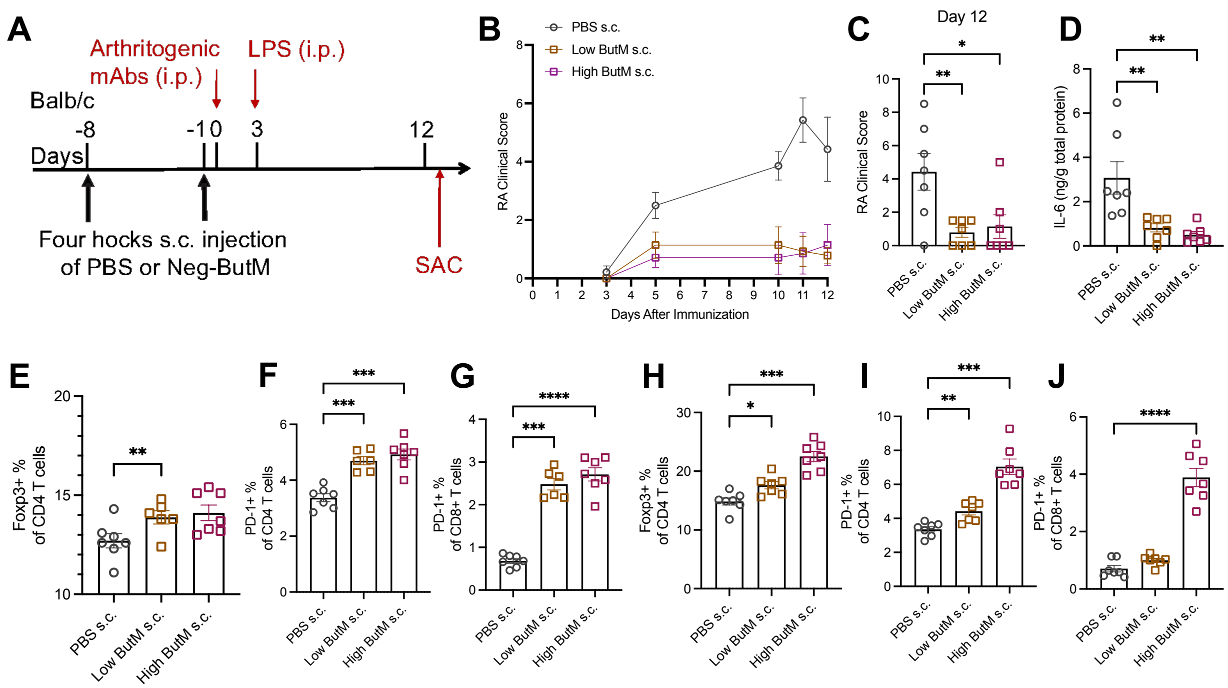


**Figure S6. Negatively charged butyrate micelles suppress arthritis progression when injected subcutaneously in four hocks. A.** Experimental scheme of the collagen antibody-induced arthritis (CAIA) model. Mice were subcutaneously injected in four hocks with PBS (*n* = 7) or Neg-ButM (2.5 or 10 mg) (*n* =7) twice on day -8 and day -1. CAIA was induced by passive immunization with an anti-collagen antibody cocktail on day 0, followed by intraperitoneal injection of lipopolysaccharide (LPS). **B.** Arthritis scores were measured daily after the immunization. The areas under the curve were compared. **C.** Arthritis scores from PBS or Neg-ButM treated mice on day 12. **D.** IL-6 measured from homogenized paws from mice treated with PBS or Neg-ButM. **E-G.** The percentage of Foxp3$^+$ regulatory T cells (E), PD-1$^+$ of CD3$^+$CD4$^+$ T cells (F), or PD-1$^+$ of CD3$^+$CD8$^+$ T cells (G) in the hock-draining lymph nodes (LNs). **H-J.** The percentage of Foxp3$^+$ regulatory T cells (H), PD-1$^+$ of CD3$^+$CD4$^+$ T cells (I), or PD-1$^+$ of CD3$^+$CD8$^+$ T cells (J) in the spleen. Experiment was repeated twice. Data are presented as mean ± SEM. Statistical analyses were performed using a one-way ANOVA with Dunnett's test. *p<0.05, **p<0.01, ***p<0.001, ****p<0.0001.

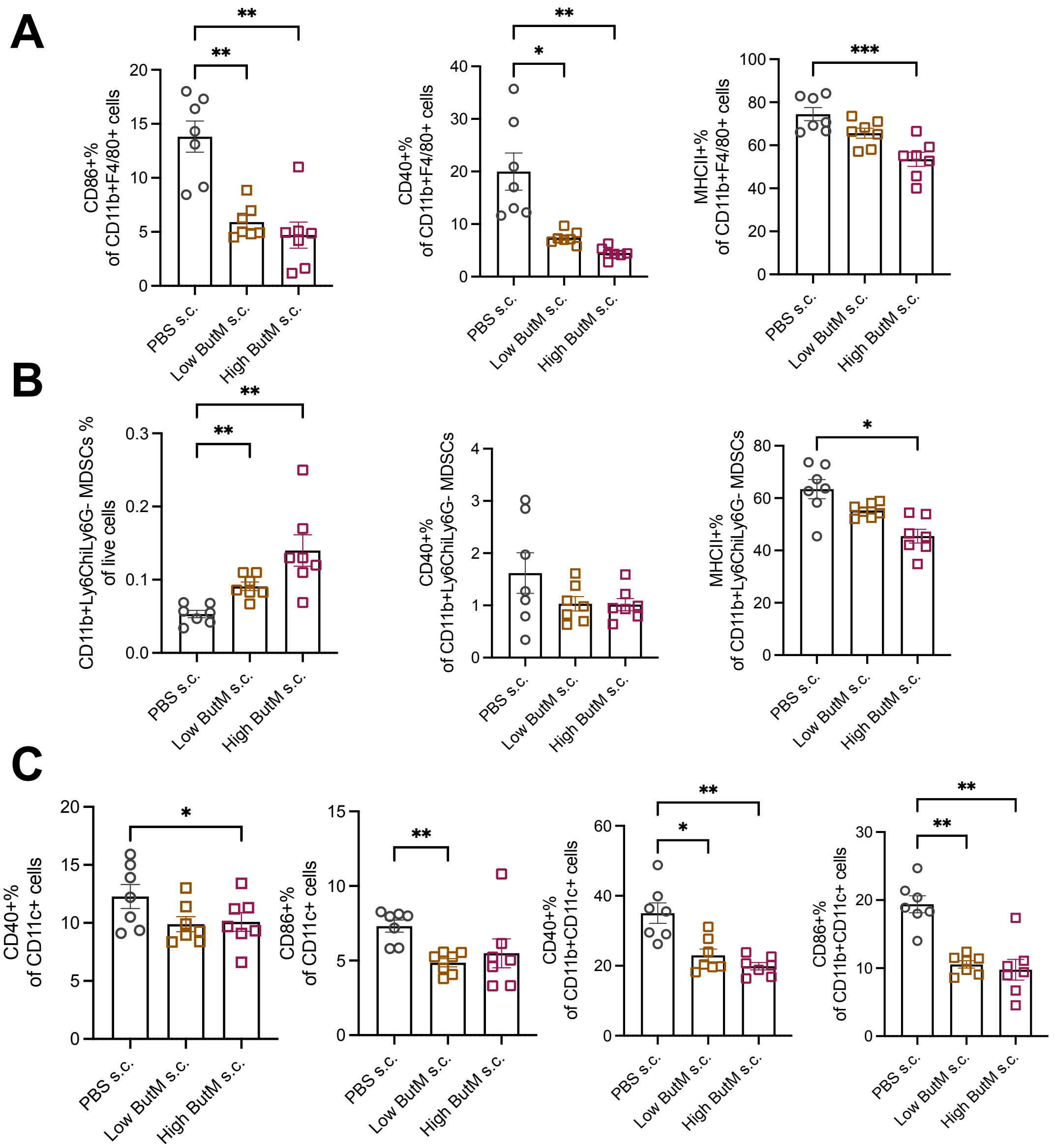


**Figure S7. Negatively charged butyrate micelles suppress myeloid cells in arthritis when injected subcutaneously in four hocks. A.** The percentage of $CD86^+$, $CD40^+$, and $MHCII^+$ of $CD11b^+F4/80^+$ macrophages in the hock-draining lymph nodes (LNs). **B.** The percentage of $CD11b^+Ly6C^+Ly6G^-$ myeloid-derived suppressor cells (MDSCs) of live cells, and $CD40^+$, and $MHCII^+$ of MDSCs in the hock-draining LNs. **C.** $CD40^+$ and $MHCII^+$ of $CD11c^+$ or $CD11c^+CD11b^+$ dendritic cells in the hock-draining LNs. Data are presented as mean ± SEM. Statistical analyses were performed using a one-way ANOVA with Dunnett's test. *p<0.05, **p<0.01, ***p<0.001, ****p<0.0001.

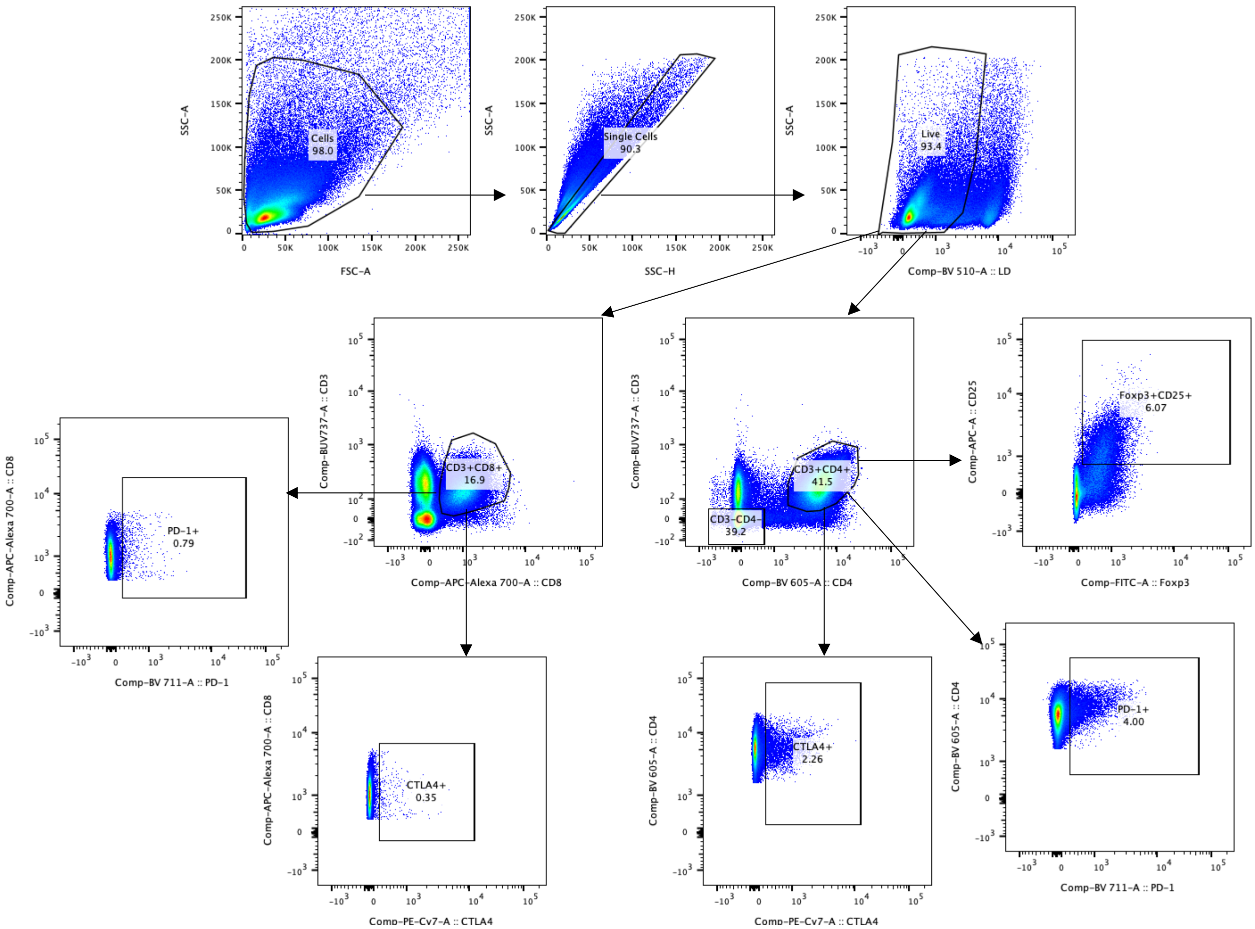


**Figure S8.** Flow cytometry gating strategy for the identification of Foxp3$^{+}$CD25$^{+}$ regulatory T cells, PD-1$^{+}$, CTLA-4$^{+}$ cells of CD3$^{+}$CD4$^{+}$ T cells, or PD-1$^{+}$, CTLA-4$^{+}$ cells of CD3$^{+}$CD8$^{+}$ T cells. This gating strategy was used in Fig.5.

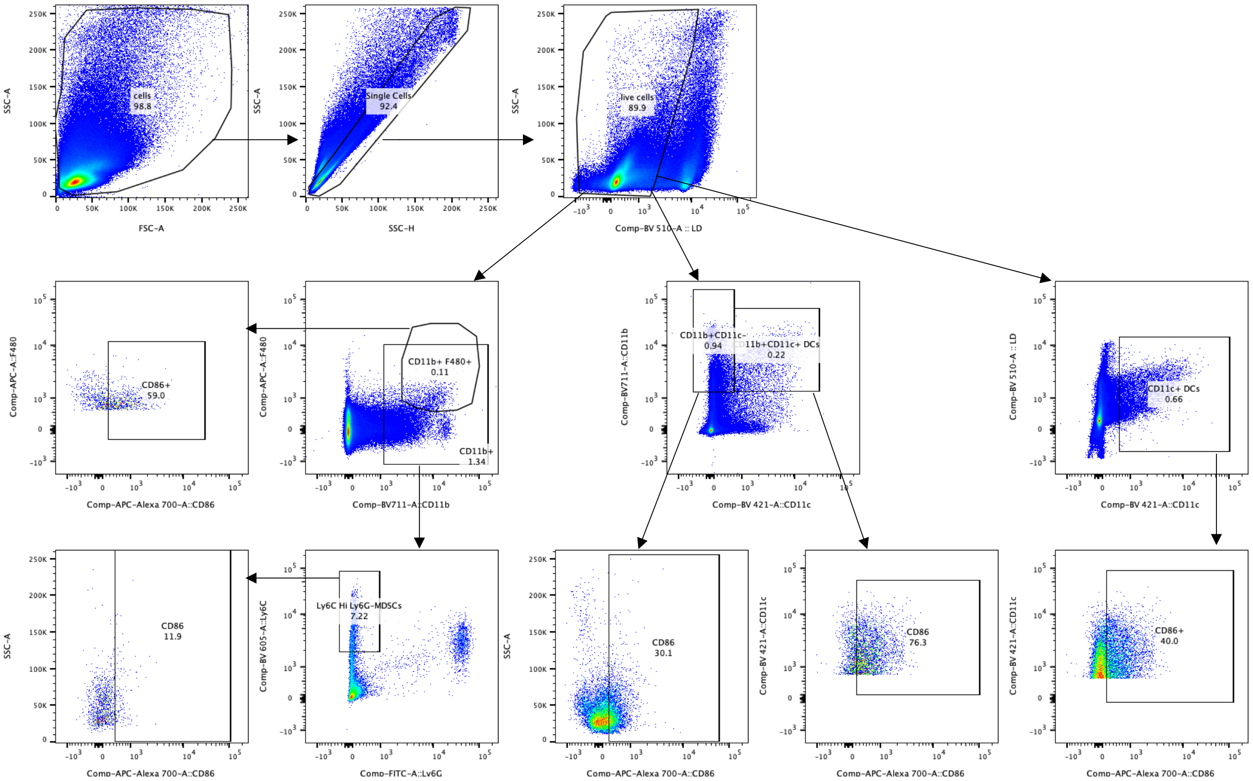


**Figure S9.** Flow cytometry gating strategy for the identification of CD86 expression in various myeloid cells, including CD11b$^+$CD11c$^-$ cells, CD11b$^+$F4/80$^+$ cells, CD11b$^+$Ly6C$^+$Ly6G$^-$ myeloid-derived suppressor cells (MDSCs), CD11c$^+$ and CD11c$^+$CD11b$^+$ dendritic cells. This gating strategy was used in Fig.6.

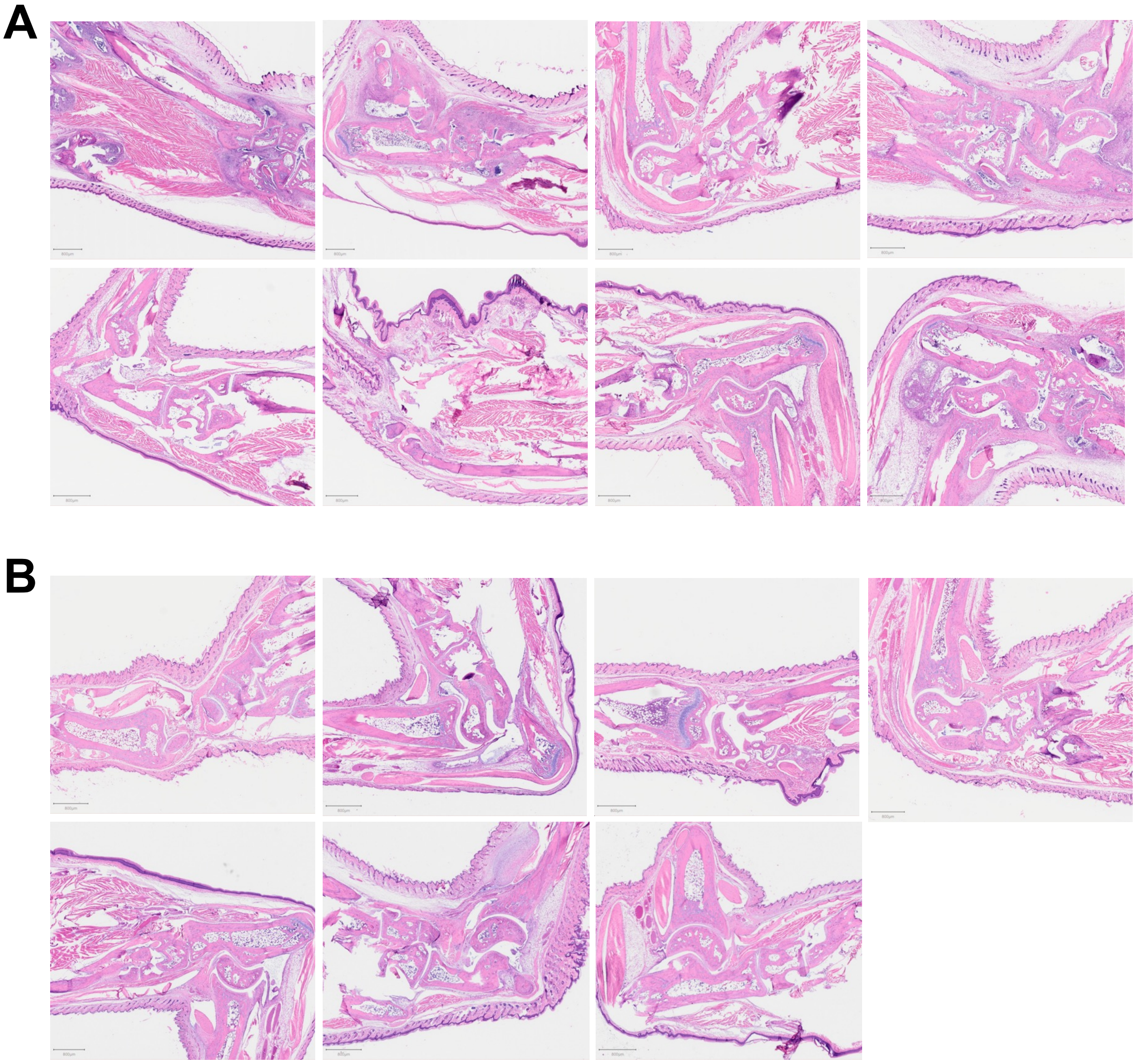


**Figure S10.** Histology images of PBS (**A**) or Neg-ButM (**B**) treated mouse hindpaws stained with hematoxylin and eosin, from the experiment in Fig.4.
